# Bivalent habenula modulation of human monoaminergic midbrain and cortical pathways

**DOI:** 10.64898/2025.12.26.696619

**Authors:** Leah J Hudson, Christopher G Davey, Trevor Steward, Po-Han Kung, Yingliang Dai, Lee Unsworth, Alec J Jamieson, Bradford A Moffat, Rebecca Glarin, Ben J Harrison

## Abstract

The habenula is implicated in signaling negative reward prediction errors (RPE), yet direct causal evidence demonstrating its influence on downstream human brain circuits is limited. Using ultra-high field (7T) fMRI and dynamic causal modelling across two negative RPE-inducing experimental tasks, we characterize habenula-directed connectivity with monoaminergic midbrain and cortical pathways in healthy adults. Negative feedback drove strong habenula inhibition of ventral tegmental area and dorsal raphe nucleus activity, while positive feedback had an excitatory effect. These core effects are replicated across reversal learning and perceptual decision uncertainty tasks, establishing a generalizable mechanism for behavioral adaptation. Task-specific modulatory effects emerged in pathways between the subcortical and cortical regions (ventral tegmental area→dorsal anterior cingulate cortex, dorsal raphe nucleus→medial prefrontal cortex, and dorsal anterior cingulate cortex→habenula), revealing context-dependent cortical elaboration of negative RPE-driven habenula signals. This provides the first evidence in humans for bivalent causal modulation of habenula function on monoaminergic midbrain and cortical pathways, demonstrating how the brain integrates negative and positive signals to guide adaptive behavior.

## Introduction

The computational encoding of reward and punishment is a fundamental neurocognitive mechanism underlying adaptive behavior^1^. Central to this process is ‘reward prediction error’ (RPE)^1^ – the difference between anticipated and received outcomes. RPE operates through a dual signaling mechanism whereby positive RPE – instances where outcomes exceed expectations – facilitate behavioral reinforcement and action repetition. Conversely, negative RPE, arising when outcomes fail to meet or oppose predictions, promotes behavioral change^2^. When functioning optimally, RPE computations enable the dynamic correction of behaviors, while serving as the motivational driver for beneficial actions^3, 4^. Although the neural circuitry underlying positive RPE has been extensively characterised^5^, the mechanisms encoding negative RPE are much less understood^6, 7^. This represents an important knowledge gap, especially considering the hypothesized role of negative RPE processes in the pathophysiology and treatment of common mental health disorders such as depression^8^.

One brain structure that has emerged as a major candidate in the neural circuitry of negative RPE is the habenula – a region that has been referred to as the brain’s ‘anti-reward centre’^2, 8, 9^. The habenula, which contains mostly glutamatergic neurons, is a highly conserved epithalamic nucleus that is extensively integrated with dopaminergic and serotonergic neurocircuitry^2, 8^. Animal studies have shown that habenula activity is generally bivalently modulated in response to aversive and appetitive feedback: aversive stimuli, including unexpected punishment (negative RPE), increases habenula activity, whereas appetitive outcomes suppress it^10–14^. This bivalent modulation exerts opposing influences on the habenula’s primary downstream targets: the dopaminergic ventral tegmental area (VTA) and serotonergic dorsal raphe nucleus (DRN)^7^. Specifically, habenula response to aversive stimuli inhibits VTA and DRN activity, while habenula suppression by appetitive stimuli increases their activity, thus indicating the habenula to be a critical orchestrator of opposing influences on these systems^10–12, 14–17^. The VTA and DRN subsequently project to higher-order regions, including the medial prefrontal cortex (mPFC) and dorsal anterior cingulate cortex (dACC)^16, 17^. Through these extended pathways, the habenula’s bivalent modulation of VTA and DRN activity decreases approach related behaviors during negative outcomes and facilitates them during positive outcomes, thereby enabling flexible, context-appropriate behavioral adaptation^16–18^.

While animal research has established a clear bivalent model of habenula function with defined anatomical pathways, human neuroimaging evidence remains incomplete^19–22^. To date, individual studies have confirmed only one component of the bivalent response; for instance, Lawson and colleagues^20^ reported increased habenula activation to aversive (conditioned threat) but not significant habenula deactivation to rewarding feedback.

Whereas Weidacker et al.^23^ observed greater habenula deactivation to positive versus negative feedback in a monetary loss avoidance task. Moreover, although these studies have reported habenula involvement alongside monoaminergic midbrain and prefrontal regions^19–22^, none have examined the influence of habenula activity changes on this extended circuitry – a critical gap given that animal studies have shown prominent direct influences of habenula projections on VTA and DRN activity^16, 17^.

To address these questions, we combined ultra-high field (7T) fMRI with two experimental paradigms designed to engage the habenula and its extended circuitry via negative RPE-like and positive feedback: a reward-punishment reversal learning task^24–27^ and a perceptual decision uncertainty task^19, 22^. Testing our predictions across two distinct task contexts enabled us to address the generalizability of feedback-driven modulation of habenula circuitry, as suggested by animal studies. We first used univariate analyses to characterize differential activation of the habenula and its extended circuitry in response to negative versus positive feedback trials across both tasks. We then employed dynamic causal modelling (DCM) to test the directional influence of feedback-driven changes in habenula activity on monoaminergic midbrain and prefrontal regions. Specifically, we hypothesized that the effect of negative feedback on habenula function would suppress VTA and DRN activity, whereas the effect of positive feedback on habenula function would enhance their activity, with these influences further propagating to the dACC and mPFC. As an extended aim, we investigated whether negative RPE encoding across the extended habenula circuitry differed as a function of specific task demands.

## Results

### Task Performance

A detailed description of the reversal learning and perceptual decision uncertainty tasks is provided in the Methods. In the reversal learning task, participants learned binary stimulus-reward associations that reversed periodically without warning. Reversals occurred only after participants demonstrated clear learning (3-6 consecutively correct trials) resulting in a mean of 27.8 ± 4.9 reversals per participant. This created conditions of unexpected punishment when previously rewarded stimuli suddenly delivered punishment; and expected reward during stable periods between reversals.

In the perceptual decision uncertainty task, participants predicted the direction of a moving dot array under high perceptual decision uncertainty and received trial-by-trial feedback of incorrect and correct responses. Task difficulty was adaptively calibrated every ten trials to ensure that participants experienced consistent prediction uncertainty during the task (mean error rate = 29%).

### Task-Based Activation and Deactivation Mapping

We first performed univariate general linear model (GLM) analyses to identify brain regions demonstrating differential activation and deactivation in response to negative versus positive feedback (false discovery rate corrected statistical threshold of *P*_­_< .05, cluster-extent threshold of *K_­_*≥ 5 voxels). Whereby negative feedback consists of unexpected punishment and incorrect response trials; positive feedback refers to expected punishment and correct response trials for the reversal learning and perceptual decision uncertainty tasks, respectively. This analysis primarily served to localize volumes-of-interest (VOIs) for subsequent DCM analyses.

Across both tasks, it was confirmed that negative > positive feedback was associated with significant activation of the habenula, VTA, DRN, and dACC (Fig. 1). Additional activation was observed bilaterally in the anterior insula cortex, dorsolateral prefrontal and premotor cortices, superior parietal cortex, caudate nucleus, and putamen.

**Figure 1.**
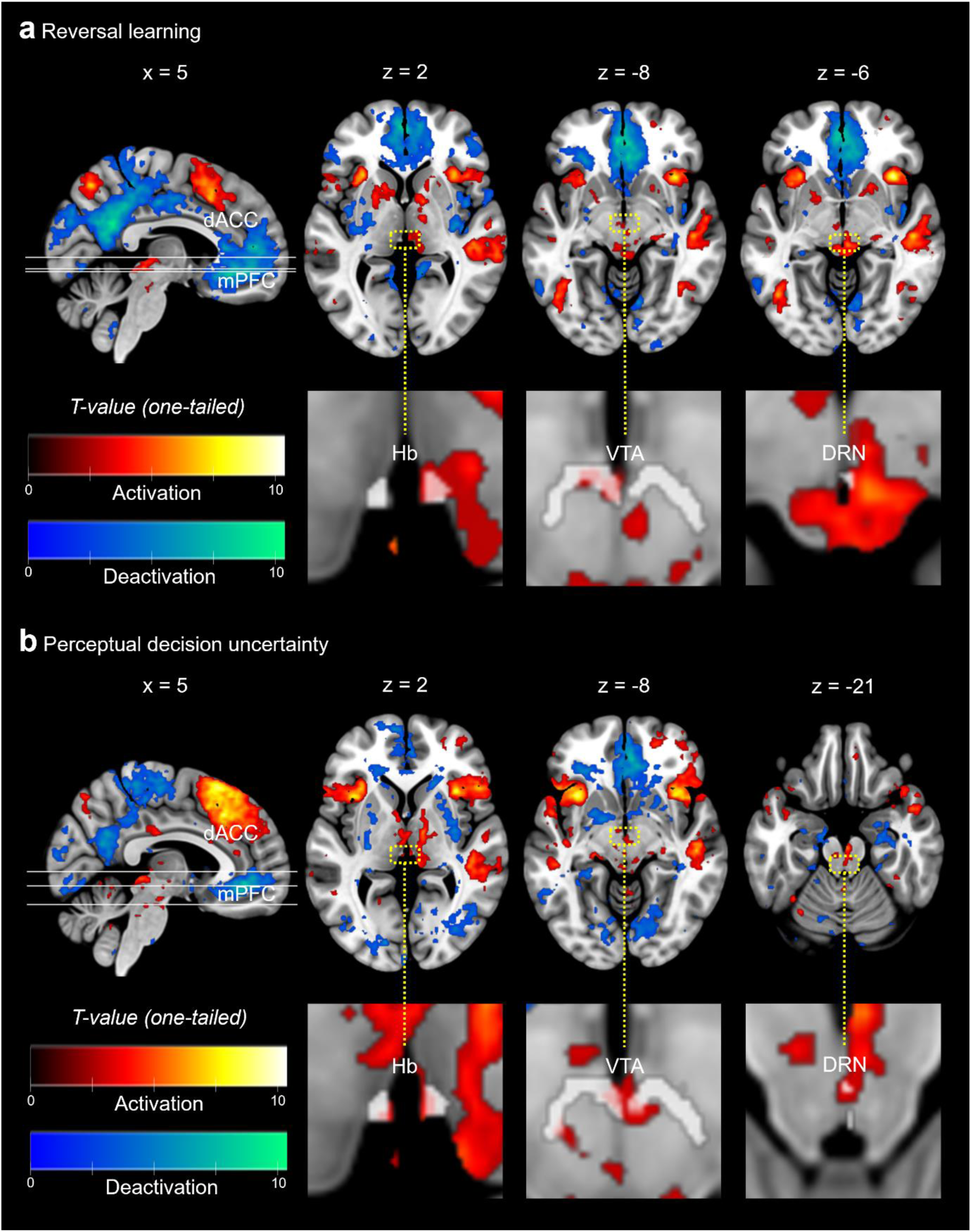
Task-based activation and deactivation. **a** Whole-brain GLM results for the reversal learning task. **b** Whole-brain GLM results for the perceptual decision uncertainty task. For both tasks, significant results (*P_­_*< .05, *K_­_*≥ 5) are presented on the MNI 152 template brain. The color maps represent corresponding *t*-statistic values for a one-tailed *t* test. Warm/cool colors correspond to regions of significant activation/deactivation during negative feedback (unexpected punishment; incorrect response) compared to positive feedback (expected reward; correct response). Our targeted regions-of-interest are highlighted in their respective anatomical planes with translucent masks outlining the subcortical regions. *dACC*, dorsal anterior cingulate cortex; *mPFC*, medial prefrontal cortex; *Hb*, habenula; *VTA*, ventral tegmental area; *DRN*, dorsal raphe nucleus; *FDR*, false discovery rate; *K_­_*, voxel cluster-extent threshold; *MNI,* Montreal Neurological Institute.

Across both tasks, negative < positive feedback was associated with significant deactivation of the mPFC (Fig. 1). Additional regions deactivated included the posterior cingulate cortex, precuneus and lateral orbitofrontal cortex. As well as the posterior insular cortex, inferior parietal cortex, and amygdala-hippocampus. Presentation of overlapping activation across both tasks can be seen in Supplementary Fig. 1. Complete GLM results are provided in Supplementary Table 1 and 2.

### Dynamic Causal Modelling of Habenula Network Interactions

Having confirmed consistent activation of the habenula and extended regions across both tasks, we used DCM to test the hypothesized bivalent influence of feedback-driven changes in habenula activity on the extended midbrain and cortical regions-of-interest (ROIs). DCM estimates ‘effective connectivity’ – the directed influence that one brain region exerts on another – through neurobiologically informed generative models^28–30^. Under a Bayesian framework, the winning model balances data fit against model complexity. The resulting effective connectivity estimates, reported in hertz (Hz), specify the rate of change in one region’s neuronal activity caused by another region’s activity. Connection strengths can be excitatory (positive values indicating activity upregulation) or inhibitory (negative values indicating activity downregulation) and can be intrinsic (baseline connectivity) or modulated by experimental conditions.

We formulated DCM models consisting of the habenula (Hb), VTA, DRN, dACC, and mPFC and tested how feedback modulated connectivity across this network (see Methods; Fig. 2). Critically, the model architecture reflects known anatomical constraints: bidirectional connections linked the subcortical regions (Hb↔VTA, Hb↔DRN, VTA↔dACC, DRN↔mPFC), but cortical regions were only unidirectionally connected to the habenula (mPFC→Hb, dACC→Hb), consistent with the absence of direct habenula-to-cortex projections (Supplementary Fig. 2). All regions included self-inhibitory connections.

**Figure 2.**
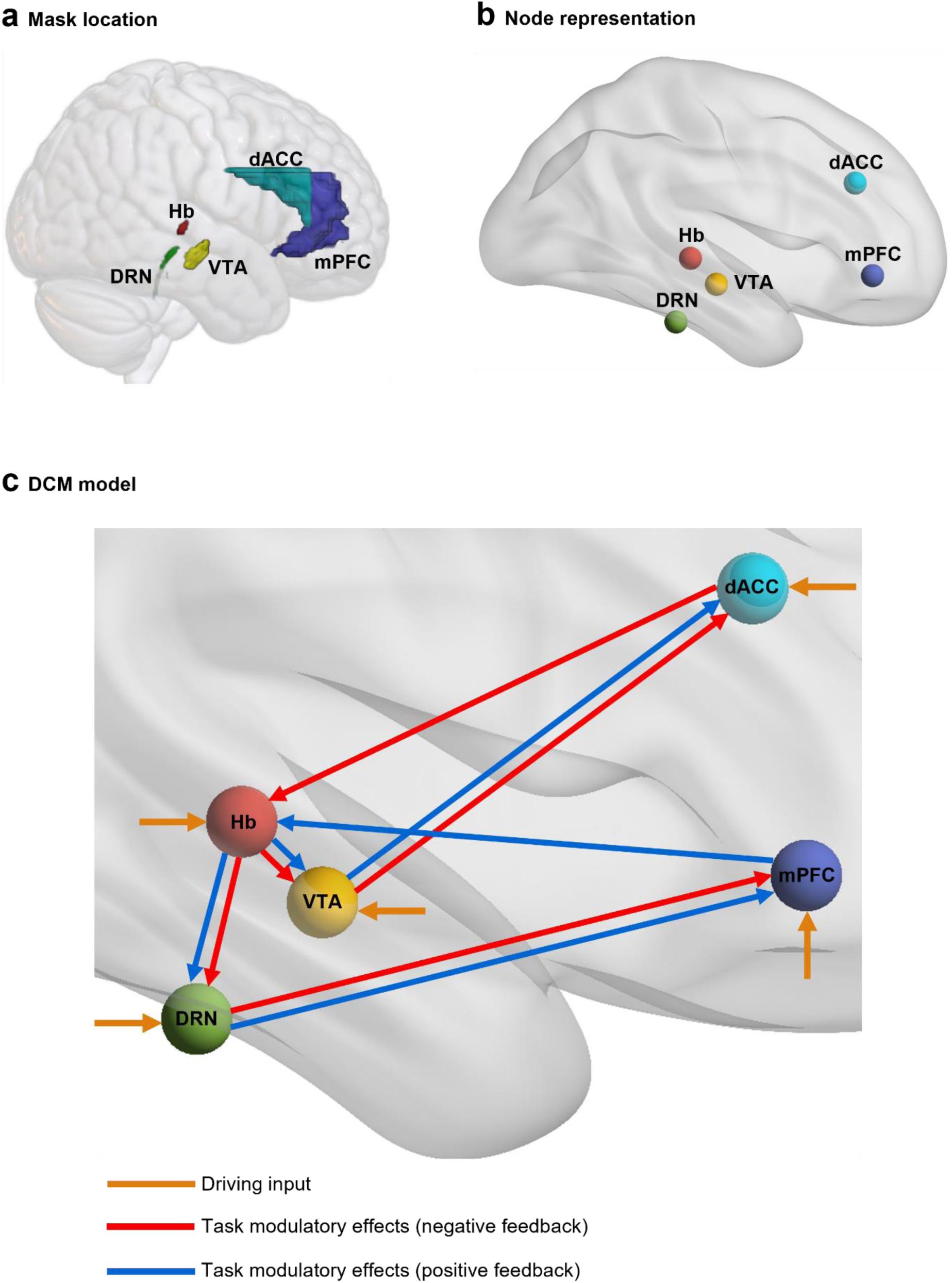
Representation of DCM model space. **a** Mask location. These five masks cover the Hb, VTA, DRN, dACC and mPFC, and informed volume-of-interest extraction for the DCM model. **b** Node representation. These nodes are for visualization purposes and represent our regions-of-interest included in the model. **c** Model architecture. The model assumed bidirectional endogenous connections between the Hb↔VTA, Hb↔DRN, dACC↔VTA, mPFC↔DRN as well as unidirectional endogenous connections from the mPFC→Hb and dACC→Hb – not pictured. Endogenous self-inhibitory connections were also modelled – not pictured. Modelled task modulatory effects are presented in red for negative feedback and in blue for positive feedback modulation. Driving input from the experimental conditions of the tasks into all regions was assumed (orange arrows). 3-D brain rendering was constructed in MRIcroGL^32^ for Fig. 2a and BrainNet Viewer^33^ for Fig. 2b and 2c with the MNI152 template brain. *DCM*, dynamic causal modelling; *GLM*, general linear modelling; *Hb*, habenula; *VTA*, ventral tegmental area; *DRN*, dorsal raphe nucleus; *dACC*, dorsal anterior cingulate cortex; *mPFC*, medial prefrontal cortex; *MNI,* Montreal Neurological Institute.

Feedback presentation (unexpected punishment and expected reward in reversal learning; incorrect and correct responses in the perceptual decision uncertainty task) served as the driving input to all regions. We assessed modulatory effects of negative feedback (unexpected punishment; incorrect response) and positive feedback (expected reward; correct response) on key connections within this network. To establish generalizability, we tested these effects across both tasks separately. Group-level connectivity estimates were obtained using Parametric Empirical Bayes (PEB)^31^ and thresholded at posterior probability (Pp) > .95.

#### Bivalent Modulation of Habenula→Midbrain Connectivity

Consistent with our primary hypothesis, negative and positive feedback bivalently modulated habenula influences on the VTA and DRN across both tasks (Fig. 3). Negative feedback drove strong habenula inhibition of VTA (reversal learning: -3.65 Hz; perceptual decision uncertainty: -1.24 Hz) and DRN activity (reversal learning: -1.61 Hz; perceptual decision uncertainty: -0.86 Hz). Conversely, positive feedback drove strong habenula excitation of VTA (reversal learning: 3.93 Hz; perceptual decision uncertainty: 2.70 Hz) and DRN activity (reversal learning: 2.72 Hz; perceptual decision uncertainty: 1.16 Hz). The modulatory effects of positive and negative feedback on habenula connectivity were consistent across both paradigms, but larger in the reversal learning task.

**Figure 3.**
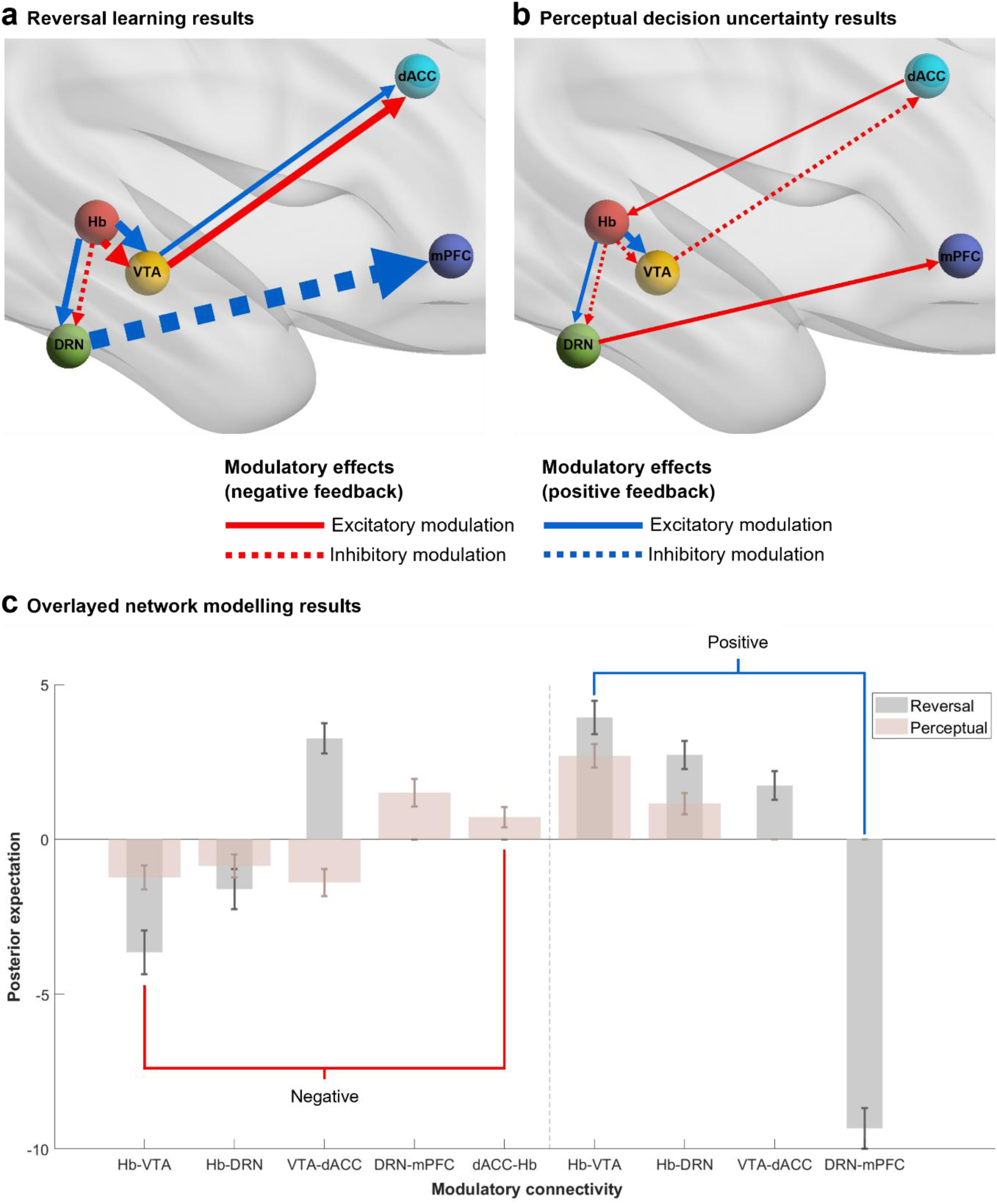
Network Modelling Results. Task-induced modulation of connections that demonstrated strong evidence (posterior probability >.95) for a non-zero group effect. Modulated connections are presented with dashed lines where respective line thickness indicates connection strength in Hz. Modulatory effects due to negative feedback are shown in red, whereas positive feedback is presented in blue. Excitatory modulation is represented with complete lines, and inhibitory modulation is represented with dashed lines. **a** Reversal learning task results. **b** Perceptual decision uncertainty task results. Illustrated using BrainNet Viewer with the MNI152 template brain^33^. **C** Comparison of the modulated network connectivity across both tasks. Each bar represents the Bayesian model-averaged (BMA) connectivity strengths as measured in Hz for each modulated connection. The reversal learning task is presented in grey with the perceptual decision uncertainty task overlayed in pink. Whiskers represent the 95% confidence interval as derived from the posterior covariance matrix (spm_plot_ci). The graph presents the network connectivity results as modulated by negative feedback (red) and positive feedback (blue). *Hb*, habenula; *VTA*, ventral tegmental area; *DRN*, dorsal raphe nucleus; *dACC*, dorsal anterior cingulate cortex; *mPFC*, medial prefrontal cortex; *MNI,* Montreal Neurological Institute.

#### VTA→dACC Pathway

Feedback modulation of midbrain-cortical pathways demonstrated greater task specificity. During reversal learning, VTA activity exerted an excitatory influence on the dACC under both negative (3.26 Hz) and positive feedback (1.73 Hz). Whereas during perceptual decision uncertainty, negative feedback drove VTA inhibition of dACC activity (-1.38 Hz), while positive feedback had no meaningful modulatory effect (-0.44 Hz; Pp > .75). These results suggests that VTA→dACC connectivity is differentially modulated by feedback context – showing a consistent excitatory influence when negative feedback signals unexpected contingency violations but inhibition when it occurs under sustained uncertainty.

#### DRN→mPFC Pathway

Task-specific modulation was also evident in the DRN→mPFC pathway. During reversal learning, positive feedback drove strong DRN inhibition of mPFC activity (-9.33 Hz), with negative feedback showing no meaningful modulation (0.68 Hz; Pp > .75). Conversely, during perceptual decision uncertainty, negative feedback drove DRN excitation of mPFC activity (1.49 Hz), while positive feedback had no detectable effect. These opposing results across tasks indicate that the propagation of habenula-driven modulation to cortical regions is context-dependent, potentially reflecting different computational demands between unexpected contingency learning and prediction under uncertainty.

#### Cortical Modulation of Habenula Activity

Only the dACC→Hb pathway showed meaningful feedback modulation, and this only occurred during the perceptual decision uncertainty task. Negative feedback drove dACC excitation of habenula activity (0.71 Hz), whereas no meaningful effect was observed during reversal learning (0.30 Hz; Pp > .5). This suggests top-down cortical regulation of habenula activity may be specific to contexts involving sustained prediction uncertainty.

Complete parameter estimates derived from Bayesian model averaging, including posterior expectation, posterior covariance and posterior probability, are reported in Supplementary Table 3.

## Discussion

This study provides the first evidence in humans for bivalent causal modulation of habenula function on midbrain monoaminergic and cortical pathways by negative and positive feedback. We have shown that negative feedback drives habenula inhibition of dopaminergic and serotonergic midbrain nuclei while positive feedback produces the opposite effect. This key finding establishes a neurobiological mechanism for how the brain dynamically shifts between behavioral persistence and adaptive change in response to outcomes. Importantly, these core effects replicated robustly across two task contexts, demonstrating that habenula-midbrain modulation operates as a generalizable property of feedback processing, independent of whether negative feedback signals unexpected contingency violations or sustained prediction uncertainty. This bivalent mechanism likely provides a neural substrate for flexible context-appropriate behavioral adaptation, with task-specific propagation to cortical regions indicating how subcortical feedback signals are integrated with higher cognitive control systems^34^. Our results extend animal findings^10–17^ to humans and establish the habenula as a critical orchestrator of opposing influences on the brain’s primary monoaminergic systems.

Across both tasks, our univariate GLM analyses confirmed consistent habenula responses to negative versus positive feedback, strongly supporting its role in encoding negative RPE across different feedback contexts – from unexpected contingency violations to sustained prediction uncertainty. These findings align with previous human neuroimaging studies^19–22^ that have reported habenula responses to aversive outcomes, while extending them by demonstrating robustness across multiple paradigms. Unlike Weidacker and colleagues^23^, we did not observe significant habenula deactivation in response to positive feedback, possibly reflecting differences in task design (monetary loss avoidance versus the reward-punishment learning) and/or analytic approaches: in their study, habenula signal decreases were characterized against an implicit baseline condition whereas here we reviewed habenula signal change across a comparison of negative and positive feedback. Our observation of mPFC deactivation during negative versus positive feedback aligns well with established reward circuitry models, where mPFC typically shows greater relative activation to reward processing and value representation^35^. Conversely, greater dACC activation during negative compared to positive feedback is broadly consistent with accumulating evidence from fMRI reward processing studies evidencing dACC reactivity to reward evaluation^21, 22, 36,37^. The concurrent activation of VTA and DRN during negative versus positive feedback contrasts with their more typically noted reward-related responses in fMRI studies^38, 39^. However, as characterized in animal studies, both regions contain neurons with opposing response properties: reward-responsive populations (suppressed by habenula input during aversive outcomes^14, 16^) and punishment-responsive populations that increase firing to aversive stimuli^6, 40–42^. Recent accounts propose that these midbrain regions encode motivational salience independently of valence^41^ ^42^, with negative feedback potentially eliciting greater salience than positive feedback in our tasks. Thus, our univariate results likely capture the dominant influence of punishment-responsive and salience-encoding neuronal populations during negative feedback.

Using hypothesis-driven DCM models, we have observed that negative and positive feedback bivalently modulates habenula influences on the VTA and DRN, providing a strong human translation of findings accumulated from animal studies^10–12, 14–17^. This bivalent modulation – with negative feedback driving strong inhibition and positive feedback driving excitation of midbrain activity – provides a neural mechanism for dynamically regulating approach-related behavior: suppressing it when outcomes are worse than expected, and facilitating it when outcomes are favorable. The consistency of these effects across both tasks indicates that habenula modulation of subcortical activity acts as a domain-general ‘switch’ for behavioral adaptation, operating independently of whether negative feedback signals unexpected contingency violations or outcome uncertainty. Importantly, this mechanism directly shapes motivated behavior through the VTA and DRN’s established roles in reward processing^43, 44^ and evaluation of motivational salience^41, 42^. Prior research has elucidated that dual involvement of the VTA and DRN is required for optimal learning^45^ with each region contributing to reward processing in differing ways: the VTA contributes to the encoding of ‘reward memory’ i.e., whether an action previously brought about a reward, whereas the DRN aids the encoding of long-term reward value^10, 40^. The observed bivalent influence of the habenula on these regions therefore endorses its role as a key regulator of reward-punishment valuation, determining whether individuals persist with or abandon current behavioral strategies based on outcome feedback.

An important goal of our study was to evaluate the down-stream influences of the habenula-midbrain interactions on cortical function. Regarding the VTA, Lammel and colleagues^17^ showed that habenula-mediated VTA signaling to the prefrontal cortex evoked punishment-related behaviors, and thus demonstrated habenula-midbrain driven behavior adaptation.

Extending this work, we specifically modelled the downstream influence of habenula-VTA connectivity on the dACC. The dACC has a well-established role in monitoring action outcomes and driving behavioral flexibility^46^ – with our findings suggesting a close integration of these functions with motivational state information provided by the VTA. Notably, the observed direction of the VTA’s influence on the dACC differed between the two tasks.

During reversal learning, the VTA had an excitatory influence on the dACC irrespective of feedback valence; whereas during decision uncertainty, the VTA had an inhibitory influence on the dACC during negative feedback alone. Thus, the propagation of habenula-VTA influences on cortical control systems appears to adaptively reconfigure based on task features. As successful reversal learning requires flexible behavioral adaptation, increased excitation from the VTA to dACC may facilitate the enhanced cognitive control required to override previously learned associations^47^. In contrast, during sustained uncertainty, VTA inhibition of the dACC may prevent premature or maladaptive behavioral changes when outcomes remain unpredictable^46^. This flexibility suggests how subcortical feedback signals may be integrated with cortical monitoring systems to produce context-appropriate behavioral regulation^18, 46^.

Task-specific modulation was also evident in the DRN→mPFC pathway, which emerged alongside observations of DRN activation and mPFC deactivation during negative versus positive feedback in the univariate analyses. Animal studies have shown DRN stimulation to exert an inhibitory influence on the mPFC^48, 49^. During reversal learning, we observed that positive feedback led to a strong inhibitory influence of the DRN on mPFC activity. However, an opposing relationship was revealed during the perceptual decision uncertainty task, where negative feedback led to the DRN having an excitatory influence on mPFC activity. As noted, positive feedback led to the habenula exhibiting an excitatory influence on DRN activity, therefore, in combination, these findings support prior research indicating increased DRN activity to have an inhibitory influence on the mPFC^48, 49^. This bivalent coupling likely provides a mechanism for integrating serotonergic neuromodulation with prefrontal cortical regulation. Given the mPFC’s role in representing reward value, planning, and decision-making^18^, and the DRN’s involvement in evaluating long-term reward outcomes^40^, this pathway likely translates aversive and appetitive signals into adjustments of prefrontal value representations that guide future choices.

We also identified a distinct regulatory role of the dACC during the perceptual decision uncertainty task, providing the first evidence in humans for cortical modulation of habenula function during feedback processing. Previous work by Trier and colleagues^50^ demonstrated increased ACC-habenula connectivity during threat processing using psycho-physiological interaction analyses; our DCM results extend these observations by demonstrating a causal influence of the dACC and linking it specifically to negative feedback contexts involving sustained uncertainty. The dACC’s established role in accumulating outcome history and signaling behavioral change^51^ suggests that this cortical-to-habenula pathway enables cognitive regulation of aversive processing. Specifically, when prediction errors signal heightened vigilance needs – particularly under sustained uncertainty – the dACC, informed by recent performance monitoring, may calibrate habenula responsiveness. This result suggests that the habenula is not merely a bottom-up aversive signal generator, but instead is embedded within cortical-subcortical circuits that enable context-appropriate cognition regulation of adaptive behavior.

The habenula comprises medial and lateral divisions, with animal studies primarily implicating the lateral habenula in negative RPE processing^2, 7, 9^. Despite 7T MRI allowing high spatial-temporal resolution for fMRI and individualized habenula parcellation, we could not distinguish specific habenula subregional contributions. Nevertheless, our results contribute to growing evidence that 7T fMRI can successfully target habenula circuitry influences on cognition and behavior^23, 38, 50, 52^. Methodologically, we leveraged two distinct tasks – reversal learning and perceptual decision uncertainty – to present negative RPE-like and positive feedback across varying contexts, enhancing generalizability. However, alternative tasks could generate both positive and negative RPE simultaneously. For example, Lin and colleagues^38^ used a magnitude learning task to model both VTA activity to positive RPE and habenula activity to negative RPE. Due to expected reward patterns in our reversal learning task, we could not comparatively assess positive RPE. Incorporating trial-by-trial accuracy predictions across both tasks would have enabled measurement of both positive and negative RPE, potentially revealing further novel circuit-level insights and enriching our understanding of RPE neural mechanisms.

By demonstrating bivalent causal modulation of habenula-midbrain connectivity across distinct feedback contexts, this study establishes a neural mechanism for flexible behavioral adaptation in humans. The key finding – that negative feedback drives habenula inhibition of dopaminergic and serotonergic midbrain nuclei while positive feedback produces the opposite effect – provides a neurobiological explanation for how the brain dynamically shifts between behavioral persistence and adaptive change in response to outcomes. This mechanism appears to operate as a generalizable property of feedback processing, functioning independently of whether negative feedback signals unexpected contingency violations or inherent outcome uncertainty. The propagation of these effects to cortical regions reveals further how subcortical feedback-driven signals are integrated with cognitive control system to enable context-appropriate behavioral regulation. Indeed, the demonstration of top-down cortical modulation of habenula activity establishes that this structure is not simply a bottom-up aversive signal generator but operates within an integrated cortico-subcortical network. Overall, by demonstrating that the habenula orchestrates opposing influences on the brain’s primary dopaminergic and serotonergic systems – rather than serving simply as an ’anti-reward center’ – our findings advance understanding of how aversive and appetitive processing are integrated to guide adaptive behavior.

## Materials and Methods

### Participants

Sixty-six healthy adults were recruited to the study via online advertisements and were deemed eligible to participate using the following criteria: (1) aged between 18-65 years; (2) no current diagnosis or history of a mood or psychotic disorder as assessed by the Mini International Neuropsychiatric Interview (MINI)^53^; (3) spoke English competently; (4) not currently taking any psychoactive medication; (5) no contraindications to MRI such as claustrophobia, incompatible metallic implants, or pregnancy; (6) were willing to comply with MRI safety policies and were capable of following study instructions. All participants had normal or corrected-to-normal vision. Participants provided written informed consent after receiving a complete description of the study protocol and completed a single session at the Melbourne Brain Centre Imaging Unit (MBCIU, University of Melbourne, Parkville). This study was approved by the Royal Melbourne Hospital Human Research Ethics Committee (RMH HREC 2023.034).

Out of the sixty-six participants, sixty-four completed the full scanning session. Data from nineteen participants were excluded from the reversal learning task analyses; due to insufficient task proficiency (n = 5; see Reversal Learning Task), excessive head motion (n = 5; see Imaging Preprocessing and Physiological Noise Correction), incomplete data (n = 2), and failed VOI timeseries extraction (n = 7; see Model space and time-series extraction).

Data from twenty-three participants were excluded from the perceptual decision uncertainty task analyses due to excessive head motion (n = 7), acquisition error (n = 1), incomplete data collection (n = 2), and failed VOI extraction (n = 13). The final sample comprised of forty-seven (23 females; mean age of 28.9 years ± 8.7 years) and forty-three subjects (20 females; mean age of 29.4 years ± 8.9 years) for the reversal learning and perceptual decision uncertainty tasks, respectively. See Supplementary Table 3 for further demographic information.

### Reversal Learning Task

For both tasks, stimuli were presented on an LCD screen (Cambridge Research Systems, UK) and viewed via a head-coil mounted reverse mirror. Responses were recorded via a two-button-box (Lumina LS-PAIR, Cedrus; San Pedro, CA USA) placed in the participants’ right hand.

The reversal learning task (Fig. 4a) was based on those used in previously published studies^24–27^, and programmed in PsychoPy (University of Nottingham; version 2023.2.2^54^). While in the scanner, subjects were presented with two vertically adjacent stimuli – a circle and a triangle, for 1500ms followed by a 1000ms fixed delay. For each trial, one of the stimuli was highlighted by a bold outline indicating for the subject to determine whether this shape was associated with a reward or not and to express their decision via a button press. No further information was provided for the first trial and therefore participants had to guess which shape was associated with a reward. Feedback was provided after each trial for 500ms in the form of a green thumbs up accompanied by a pleasant tone for correctly assessing which shape was associated with a reward, a red thumbs down with a low tone for incorrect responses, and text that read, “Too slow!” if a button press was not registered during stimuli presentation (1500ms). Once feedback had been received, participants could learn which shape was associated with a reward. A pseudo-random learning criterion was implemented varying from 3-6 consecutively correct trials. Once this was achieved, the reward association was reversed. If participants made an incorrect selection prior to the fulfillment of the learning criterion, the criterion was reset, and a reversal would not occur until another 3-6 trials were consecutively correct. The reversal was revealed to participants either via unexpected punishment (shape previously associated with a reward now associated with punishment) or unexpected reward (shape previously associated with punishment now associated with a reward). Participants completed an average of 165 trials – each trial was followed by a fixation cross lasting between 1000-3000ms. The average number of reversals achieved was 27.8 (± 4.9 trials) – participants that completed 19 or less reversals were excluded for insufficient task proficiency. It should be noted that successful reversals were only recorded when followed by demonstrated updated learning of the reward association i.e., when a correct selection was made directly following the reversal. As an incentive, participants were informed that they would receive an additional $25 if they performed well during the task. During debriefing, participants were informed that to perform well, they only needed to start the task and that all participants that completed their participation in the study had received the additional $25.

**Figure 4.**
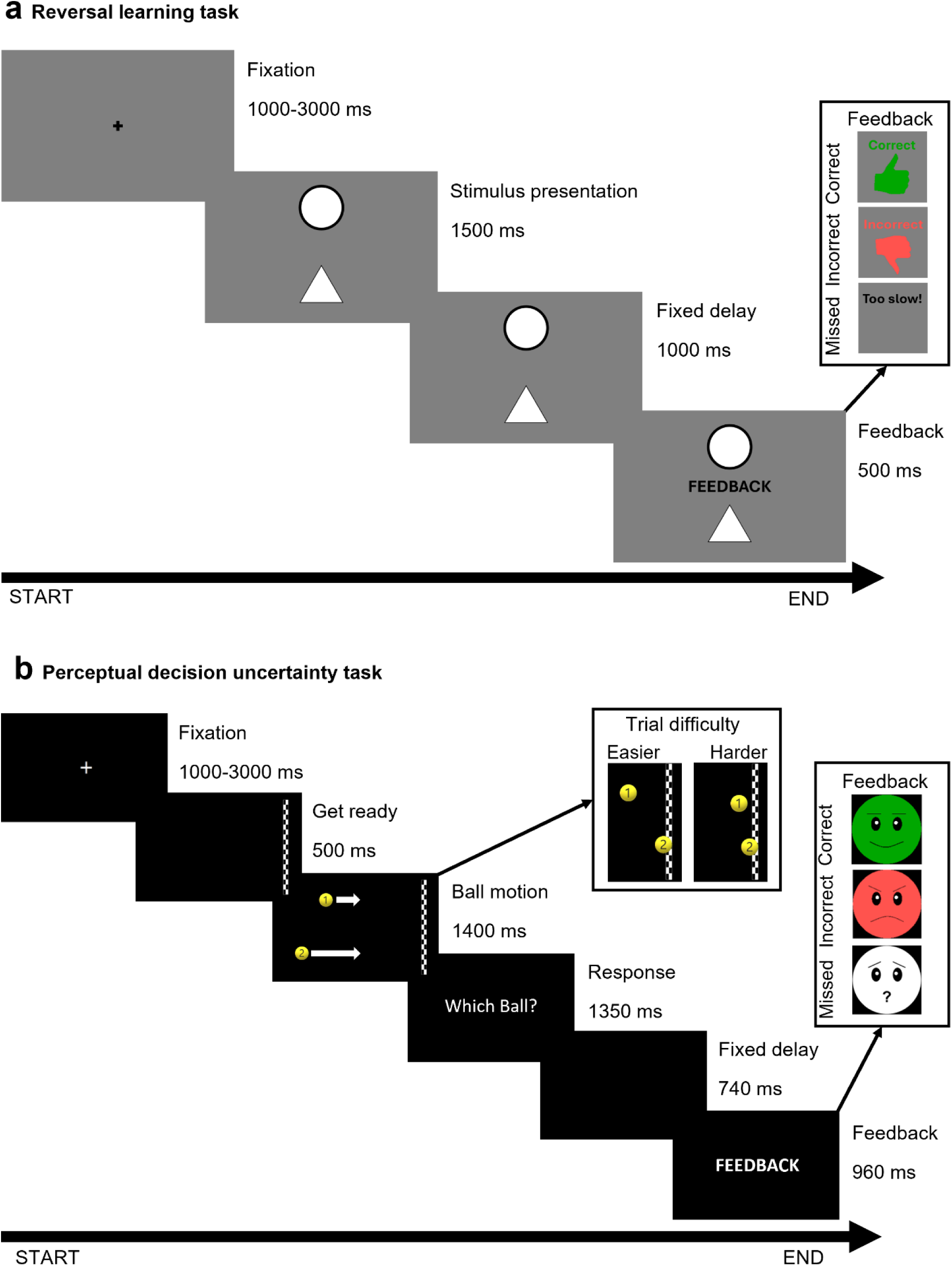
Overview of the implemented paradigms and their timings. **a** Reversal learning task. Participants were asked to learn which shape was associated with a reward. After the association was learnt, the reward association switched between the two shapes, and the participants had to re-learn which shape was associated with a reward. **b** Perceptual decision uncertainty task. Within the perceptual decision uncertainty task, participants were presented with two balls racing towards a finish line. Once the balls had disappeared, participants were asked to predict which ball would reach the finish line first. Difficulty was updated every ten trials to maintain uncertainty. Easier trials led to a larger gap between the winning and losing ball, whereas harder trials had a smaller gap – see Trial difficulty. Both tasks provided participants with informative feedback of either correct with a positive accompanying tone, incorrect with a negative accompanying tone or missed with a neutral accompanying tone (i.e., no button press recorded within the response window) – see Feedback.

### Perceptual Decision Uncertainty Task

Participants, additionally, completed an adapted version of the perceptual decision uncertainty task (Fig. 4b) developed by Ullsperger and von Cramon^19^, and Flannery and colleagues^22^. This task was programmed in E-Prime 3.0 software (Psychology Software Tools, Pittsburgh, PA) and presented to participants following the reversal learning task. During each trial, a finish line appeared on the right side of the screen for 500ms before two balls appeared on the left side of the screen and raced towards the finish line for 1400ms. The balls had variable starting points and velocities across each trial and would disappear prior to reaching the finish line. Participants were then asked to predict “Which Ball?” won the race and to indicate their choice via a button press within 1350ms. After a 740ms fixed delay, participants were presented with feedback for 960ms. Correct responses resulted in feedback that consisted of a green smiley face alongside a positive sound, incorrect responses prompted feedback that consisted of a red angry face alongside a negative tone, and trials without registered button presses resulted in a white confused face alongside a neutral tone. Between each of the 96 trials, participants were presented with a fixation cross lasting 1000-3000ms. Every 10 trials, the task difficulty was updated to encourage an error rate of 30-40%. An average error rate of 29% was achieved. Difficulty was determined by how close the balls were to each other when crossing the finish line (Fig. 4b).

### Functional Magnetic Resonance Imaging Acquisition

Imaging was performed on a Magnetom Plus 7T MRI system (Siemens Healthcare, Erlangen, Germany) equipped with a 32-channel head-coil (Nova Medical Inc., Wilmington MA, USA). The functional sequence consisted of a multi-band (factor 6) and grappa (factor 2) accelerated GE-EPI sequence in the steady state (TR = 800ms; TE = 22.2 ms; flip angle = 45°; field-of-view = 20.8 cm; slice thickness = 1.6 mm, no gap; 130 x 130-pixel matrix; 84 interleaved axial slices aligned to AC-PC line)^55, 56^. A T1-weighted high-resolution anatomical image (MP2RAGE)^57^ was acquired for each participant to assist with functional time series co-registration (TR = 5000 ms; TE = 2.04 ms; inversion times = 700/2700ms; flip angle = 4/5°; field-of-view = 24 cm; slice thickness = 0.75 mm, no gap; 330 x 330mm pixel matrix; 84 sagittal slices aligned parallel to the midline). To assist with head immobility, foam-padding inserts were placed on either side of the participant’s head. Cardiac and respiratory recordings were sampled at 50 Hz using a Siemens (Bluetooth) pulse-oximeter and respiratory belt. Information derived from these recordings was used for physiological noise correction (see Imaging Preprocessing and Physiological Noise Correction).

### Imaging Preprocessing and Physiological Noise Correction

Imaging data was pre-processed using Statistical Parametric Mapping 12 (SPM12, v7771, Wellcome Trust Centre for Neuroimaging, London, UK) within the MATLAB 2023a environment (The MathWorks Inc., Natick, MA). Motion artifacts were corrected by realigning each participant’s time series to the mean image, and resampling was completed for all images with the application of 4^th^ Degree B-Spline interpolation. Individualized motion regressors were created utilizing the Motion Fingerprint toolbox to account for movement^58^. Participants were excluded if they had a mean total scan-to-scan displacement over 1.7 mm (more than a single voxel size). Each participant’s anatomical T1 image was co-registered to the respective mean functional image, segmented and normalized to the International Consortium of Brain Mapping (ICBM) template using the unified segmentation plus Diffeomorphic Anatomical Registration Through Exponentiated Lie Algebra (DARTEL) approach^59^. Smoothing was applied with a 2mm^3^ full-width at-half-maximum (FWHM) gaussian kernel to preserve spatial specificity.

Physiological noise was modelled at the first level using the PhysIO Toolbox^60^. This toolbox applies noise correction to fMRI sequences using physiological recordings and has been found to improve blood-oxygen level dependent (BOLD) signal sensitivity and temporal signal-to-noise ratio (tSNR) at 7T^61^. The Retrospective Image-based Correction function (RETROICOR)^62^ was applied to model the periodic effects of heartbeat and breathing on BOLD signals, using acquired cardiac/respiratory phase information. The respiratory response function (RRF)^63^, convolved with respiration volume per time (RVT) was used to model low frequency signal fluctuations, which arose from changes in breathing depth and rate. Heart rate variability (HRV) was convolved with a predefined cardiac response function (CRF)^64^ to account for BOLD variances due to heart rate-dependent changes in blood oxygenation. Individualized DARTEL tissue maps segmented from each participant’s respective anatomical scan were used to apply aCompCor, which models negative BOLD signals using principal components derived from white matter and cerebrospinal fluid (CSF)^65^.

### Habenula Segmentation

As detailed in Kung et al.^52^ and replicated here, individualized habenula masks were generated for each participant. Masks were generated within the native space using participants’ high-resolution anatomical images (0.75 mm isotropic) and the Multiple Automatically Generated Templates Brain Segmentation Algorithm (MAGeTbrain)^66, 67^. This process utilized 5 habenula atlases alongside 21 of our participant’s anatomical images (acting as templates) to generate 105 candidate segmentations. These segmentations were fused to each participant’s anatomical image via a majority vote in which each voxel was assessed as either habenula or non-habenula with the most frequent label adopted in the resulting anatomical habenula mask^52, 67^.

These anatomical masks were resampled with trilinear interpolation to the mean functional image resolution (2mm isotropic) and normalized to the standard space using their corresponding DARTEL flowfields. The resulting masks were binarized at a conventional threshold of 0.2^52^. To reduce the impact of interpolation during image down-sampling and minimize non-neuronal noise arising from the CSF, an iterative volume optimization strategy was employed – see Kung et al.^52^ for further information.

### General Linear Modelling Analysis

Each participant’s pre-processed time-series and nuisance regressors (i.e., physiological noise and Motion Fingerprint regressors) were included in the first level GLM analysis. The onset and duration times were specified for each condition (primary regressor) and convolved with the SPM canonical hemodynamic response function (HRF). The implicit baseline consisted of fixation periods only. Low-frequency noise was high-pass filtered at 128-Hz and SPM’s FAST method was utilized to estimate temporal autocorrelation due to yielding superior reliability to AR(1) for short TRs^68^.

For the reversal learning task, eight conditions were modelled: expected reward (positive feedback presented following accurate learnt association of a shape with a reward), expected punishment (positive feedback presented after accurate association of a shape with punishment), unexpected reward (negative feedback presented following inaccurate association of a shape with punishment, despite this previously being true – can only occur after a reversal and if the following trial demonstrates updated learning), unexpected punishment (negative feedback presented following inaccurate association of a shape with a reward, despite previously being true – can only occur after a reversal and if the following trial demonstrates updated learning), incorrect reward (feedback presented due to the participant providing an inaccurate response, despite no reversal occurring, or failed to respond, whilst the highlighted shape was associated with reward), incorrect punishment (feedback presented due to the participant providing an inaccurate response, despite no reversal occurring, or failed to respond, whilst the highlighted shape was associated with punishment), response window, and feedback delay. Our conditions of interest were unexpected punishment (UP) and expected reward (ER) due to the negative RPE evoking nature of UP and the consistent presentation of positive feedback during ER; primary contrast images were created for UP > ER and UP < ER to categorize brain activation and deactivation in response to negative compared to positive feedback. These contrasts were included in the second-level random-effects GLM analysis (one-tailed *t*-test).

Six conditions were modelled for the perceptual decision uncertainty task: correct feedback (CorrFB; presented due to correctly predicting which ball would reach the finish line first), incorrect feedback (IncorrFB; presented due to incorrectly predicting which ball would reach the finish line first), missed feedback (presented due to no response provided during the response window), ball motion (the period in which the finish line appears and the balls race towards the finish line), which ball screen (the presentation of the response window and the following blank screen), and end of task screen. Primary contrast images were created for IncorrFB > CorrFB and IncorrFB < CorrFB to categorize brain activation and deactivation in response to negative compared to positive feedback. These contrasts were included in the second-level random-effects GLM analysis (one-tailed *t*-test).

For all GLM analyses, whole-brain, false discovery rate corrected statistical thresholds were applied (*P_­_<*.05, 5 *K_­_*). GLM activation results are provided in Fig. 1 and Supplementary Tables 1 and 2. These results informed the implemented dynamic causal model.

### Dynamic Causal Modelling

DCM estimates the directional relationships between brain regions-of-interest (i.e., effective connectivity) both intrinsically (task invariant connectivity) and modulation due to specific experimental conditions at an individual level^28^. This is accomplished by comparing different hypothetical models of how these regions may interact, and identifying the one with the most relative evidence that also minimizes model complexity. The resulting estimated parameters are indicative of the rate of change in activity of a given region as a function of the activity of another region and thus are reported in Hz. These modulations can be excitatory (positively valued upregulation of activity) or inhibitory (negatively valued downregulation of activity).

#### Model space and time-series extraction

DCM-specific GLMs were constructed for the time-series extraction, which involved the addition of the condition ‘Task’ to represent the experimental input into the model. ‘Task’ for both the reversal learning task and perceptual decision uncertainty task consisted of the onsets of the primary feedback conditions of interest i.e., ER and UP, and CorrFB and IncorrFB, respectively. Prior to VOI extraction, participant timeseries were pre-whitened to minimize serial correlations and high-pass filtered. All nuisance effects not captured by the “effects of interest” *F*-contrast were regressed out from the timeseries.

Our DCM network specified ROIs consisted of the habenula, VTA, DRN, dACC and mPFC. All ROIs were activated, except for the mPFC which was deactivated, during negative compared to positive feedback (UP > ER; IncorrFB > CorrFB). ROIs for the cortical regions (dACC and mPFC) were defined as 4mm radius spheres centered around the individual neural activation/deactivation maxima and restricted to within their respective automated anatomical labelling atlas (AAL) delineations^69^. The specific masks used were a combination of the left and right supracallosal anterior cingulate cortex masks (ACC_sup_L and ACC_sup_R) for the dACC ROI and a combination of the subgenual and pregenual anterior cingulate cortex masks for the mPFC ROI. These 4mm radius spheres were further constrained within 8mm from the group peak (Table 1).

**Table 1.** Volume-of-interest group peak coordinates in MNI space.

| Region-of-interest | dACC |  |  | mPFC |  |  |
| --- | --- | --- | --- | --- | --- | --- |
|  | x | y | z | x | y | z |
| **Reversal learning** | -8 | 22 | 26 | 2 | 46 | 10 |
| **Perceptual decision uncertainty** | 5 | 32 | 27 | 8 | 38 | -5 |

*MNI*, Montreal Neurological Institute; *dACC*, dorsal anterior cingulate cortex; *mPFC*, medial prefrontal cortex.

For all subcortical regions, masks were utilized to specify our ROIs during UP > ER or IncorrFB > CorrFB. As specified above, the combined bilateral habenula masks were generated via the MAGeT algorithm^66, 67^ as implemented by Kung and colleagues^52^. Both the VTA and DRN masks were generated using the Brainstem Navigator and consisted of the combined left and right VTA – parabrachial pigmented nucleus complex masks^70^ for our VTA ROI mask and a combination of the DRN and median raphe nucleus masks^71^ for our DRN ROI mask. These ROIs were used, alongside SPM’s principal eigenvariate function, to extract VOIs containing meaningful activation (*p <* .05, uncorrected) for the specified task contrasts at the single subject level. The statistical threshold was incrementally relaxed to *p <* .5 (uncorrected) for any ROIs that did not produce a peak coordinate with a 5-voxel cluster-extent threshold at the pre-set statistical threshold. This allowed for a minimal exclusion of participant data, and it has been noted that data lacking strong responses can still provide valuable information^29^.

#### Model specification and estimation

The model was specified using DCM 12.5 and consisted of (1) the A-matrix, i.e., the endogenous connections between and within each chosen region (habenula, VTA, DRN, dACC and mPFC; see Supplementary Figure 2), (2) the B-matrix, i.e., the modulatory effect of the task conditions on inter-region connectivity (see Fig. 2b), and (3) the C-matrix, i.e., the driving influence of the task stimuli (the ‘Task’ condition). The endogenous connections (A-matrix) were primarily bidirectional except for connections between the habenula and cortical regions due to a lack of direct habenula to cortex projections^7^. For this model, the B-matrix comprised of either UP or ER for the reversal learning analyses or of either IncorrFB or CorrFB for the perceptual decision uncertainty analyses. As previously noted, the ‘Task’ condition combined the feedback conditions of interest and this C-matrix was modelled as a driving input to all regions.

#### Parametric Empirical Bayes

The PEB framework provided group-level analysis of effective connectivity in the frame of a Bayesian hierarchical model, estimating random effects on parameters from the DCM model, including from both the A-and B-matrices. This estimation takes into account individual subject expected parameter strength and the posterior covariance or uncertainty of the estimates^31^. This PEB model investigated the between-subject commonalities across the connectivity parameters for each task separately, and thus, in the design matrix, incorporated an intercept term (single column of ones) for the overall mean connectivity. The PEB framework maximizes the chances of identifying parameter estimates typical to the group and related to task effect, whilst down-weighting noisy data and uncertain parameter estimates. Parameters from both the A- and B-matrices were summarized in this model to account for potential conditional dependency.

After the PEB model was inverted, we used Bayesian model reduction (BMR) to search and compare the relative evidence of possible reduced models, by iteratively pruning parameters that did not contribute to an increase in model evidence^72^. With the use of BMR, the optimal model structure was inferred through selecting the best accuracy-complexity balanced model. The resulting patterns of effective connectivity are illustrated in Fig. 3 at a threshold of *Pp >* .95, indicating parameters which had a *strong* level of evidence.

## Data availability

Deidentified network connectivity data for this study are publicly available at https://github.com/leahjordanhudson/Bivalent_Hb_DCM.

## Code availability

Scripts used to generate the main results and figures of this study are available at https://github.com/leahjordanhudson/Bivalent_Hb_DCM.

## Acknowledgements

We thank Ali Stevens, Tudor Sava, Ariel Kim, Paula Faimann, Holly Carey, and Xiaohui Wu for their contribution to data collection. We acknowledge the technical and scientific assistance of the Australian National Imaging Facility – a National Collaborative Research Infrastructure Strategy (NCRIS) capacity at the Melbourne Brain Centre Imaging Unit (MBCIU), The University of Melbourne. The multiband fMRI sequence was generously supported by a research collaboration agreement with CMRR, The University of Minnesota. Siemens Healthineers (Germany) provided the MP2RAGE sequence. This study was supported by the National Health and Medical Research Council of Australia (NHMRC) Ideas Grant (2027662) to B.J.H.

## Author contributions

L.J.H., C.G.D., and B.J.H. conceived the general concept of this study, designed the experiment, and developed the model with input from T.S., A.J.J., and P-H.K.; L.J.H., Y.D., L.U., A.J.J., B.A.M., R.K.G., and B.J.H. aided with data collection and the crafting of the imaging protocol. L.J.H. conducted the data analysis with support from A.J.J, P-H.K., Y.D., L.U., and B.J.H; C.D.G and B.J.H. provided supervision throughout the study. L.J.H and B.J.H wrote the original draft of the manuscript. All authors reviewed and approved the final edit of this manuscript.

## Correspondence

All correspondence and request for materials should be addressed to Ben J Harrison.

