## Supplementary Information for "Bivalent habenula modulation of human monoaminergic midbrain and cortical pathways"

### Supplementary Tables

**Supplementary Table 1. Significant reversal learning task-induced activation.**

| | MNI coordinates | | | Statistics | |
|---|---|---|---|---|---|
| | x | y | z | $K_E$ | T value |
| **UP > ER** | | | | | |
| Insula, R | 35 | 19 | -6 | 17153 | 10.14 |
| Inferior parietal gyrus, excluding supramarginal and angular gyri, L[a] | -32 | -43 | 38 | 10185 | 9.64 |
| Superior temporal gyrus, R | 50 | -22 | -6 | 2841 | 7.49 |
| Lenticular nucleus, putamen, L | -19 | 5 | -2 | 499 | 6.91 |
| Thalamus, pulvinar medial, R[a] | 0 | -30 | 2 | 872 | 6.72 |
| Middle temporal gyrus, L | -45 | -62 | -5 | 466 | 6.67 |
| Lobule VI of cerebellar hemisphere, L | -30 | -58 | -32 | 70 | 5.60 |
| Lenticular nucleus, pallidum, R[a] | 13 | 2 | 2 | 360 | 5.51 |
| Fusiform gyrus, R[a] | 38 | -58 | -8 | 115 | 5.44 |
| Lobule VI of cerebellar hemisphere, R | 30 | -62 | -21 | 159 | 5.29 |
| Hippocampus, L[a] | -21 | -43 | 5 | 19 | 4.90 |
| Red nucleus, R[a] | 2 | -13 | -10 | 37 | 4.74 |
| Calcarine fissure and surrounding cortex, R | 24 | -58 | 8 | 45 | 4.63 |
| Lobule III of cerebellar hemisphere, L[a] | -5 | -30 | -14 | 18 | 4.62 |
| Middle frontal gyrus, L | -38 | 54 | 19 | 215 | 4.56 |
| Fusiform gyrus, R | 37 | -50 | -14 | 24 | 4.46 |
| Middle temporal gyrus, L | -51 | -30 | -3 | 104 | 4.46 |
| Anterior orbital gyrus, R | 24 | 53 | -16 | 51 | 4.44 |
| Thalamus, lateral geniculate, R | 22 | -24 | -6 | 26 | 4.43 |
| Lobule III of cerebellar hemisphere, L[a] | -3 | -40 | -29 | 9 | 4.40 |
| Crus II of cerebellar hemisphere, L | -3 | -82 | -34 | 41 | 4.39 |
| Superior temporal gyrus, L | -48 | -21 | 8 | 151 | 4.35 |
| Temporal pole: middle temporal gyrus, R | 48 | 8 | -27 | 8 | 4.30 |
| Lobule VI of cerebellar hemisphere, L | -29 | -46 | -26 | 10 | 4.29 |
| Crus I of cerebellar hemisphere, L | -37 | -46 | -32 | 28 | 4.28 |
| Red nucleus, R[a] | 8 | -24 | -16 | 33 | 4.27 |
| Caudate nucleus, L | -13 | -6 | 19 | 8 | 4.27 |
| Thalamus, pulvinar medial, R[a] | 18 | -32 | 19 | 17 | 4.26 |
| Inferior temporal gyrus, R | 48 | -66 | -6 | 12 | 4.14 |
| Temporal pole: middle temporal gyrus, R | 53 | 2 | -19 | 13 | 4.09 |
| Middle temporal gyrus, L | -69 | -32 | 0 | 9 | 4.06 |
| Superior temporal gyrus, L[a] | -38 | -45 | 18 | 11 | 3.99 |
| Lobule VI of cerebellar hemisphere, L | -10 | -69 | -27 | 50 | 3.98 |
| Superior frontal gyrus, dorsolateral, R[a] | 32 | 66 | 10 | 17 | 3.97 |
| Superior frontal gyrus, dorsolateral, R | 30 | 62 | -3 | 22 | 3.97 |
| Lobule VI of cerebellar hemisphere, R | 32 | -51 | -32 | 13 | 3.94 |
| Caudate nucleus, R | 16 | 8 | 16 | 7 | 3.87 |
| Inferior temporal gyrus, L | -40 | -45 | -19 | 8 | 3.86 |
| Unknown[a] | -2 | -26 | -40 | 6 | 3.85 |
| Thalamus, ventral posterolateral, R | 21 | -26 | 6 | 7 | 3.84 |
| Supramarginal gyrus, R | 46 | -43 | 37 | 30 | 3.82 |

| | | | | | |
|---|---|---|---|---|---|
| Caudate nucleus, L[a] | -3 | 3 | 14 | 16 | 3.79 |
| Middle temporal gyrus, L | -67 | -35 | 8 | 18 | 3.79 |
| Caudate nucleus, R[a] | 6 | 0 | 18 | 17 | 3.77 |
| Anterior cingulate cortex, supracallosal, L | -8 | 24 | 24 | 10 | 3.76 |
| Middle frontal gyrus, R | 30 | 53 | 35 | 7 | 3.75 |
| Precuneus, L | -21 | -51 | 3 | 7 | 3.74 |
| Crus I of cerebellar hemisphere, L | -37 | -66 | -30 | 5 | 3.74 |
| Lenticular nucleus, pallidum, L | -11 | -3 | -2 | 6 | 3.70 |
| Middle temporal gyrus, L | -62 | -34 | 2 | 10 | 3.69 |
| Middle temporal gyrus, R | 64 | -32 | -10 | 6 | 3.68 |
| Hippocampus, L[a] | -18 | -38 | 14 | 18 | 3.66 |
| Middle occipital gyrus, L | -35 | -86 | 14 | 12 | 3.66 |
| Superior frontal gyrus, dorsolateral, R | 32 | 59 | 18 | 61 | 3.65 |
| Lobule VIII of cerebellar hemisphere, R | 19 | -67 | -51 | 6 | 3.63 |
| Lobule VI of cerebellar hemisphere, R | 37 | -53 | -27 | 12 | 3.61 |
| Superior parietal gyrus, L | -16 | -82 | 53 | 11 | 3.60 |
| Thalamus, pulvinar medial, L[a] | -11 | -27 | 14 | 15 | 3.59 |
| Fusiform gyrus, R | 32 | -50 | -19 | 13 | 3.56 |
| Middle frontal gyrus, L | -46 | 50 | 13 | 7 | 3.53 |
| Lobule VI of cerebellar hemisphere, R | 8 | -72 | -22 | 22 | 3.53 |
| Precentral gyrus, R | 58 | 3 | 43 | 9 | 3.53 |
| Temporal pole: superior temporal gyrus, L | -58 | 8 | -11 | 8 | 3.50 |
| Middle cingulate & paracingulate gyri, R[a] | 14 | -24 | 27 | 7 | 3.49 |
| Superior temporal gyrus, L | -67 | -21 | 5 | 10 | 3.47 |
| Crus I of cerebellar hemisphere, L | -32 | -74 | -24 | 7 | 3.47 |
| Fusiform gyrus, L | -30 | -51 | -21 | 8 | 3.46 |
| Calcarine fissure and surrounding cortex, L | -22 | -56 | 5 | 8 | 3.44 |
| Cuneus, R | 19 | -64 | 32 | 5 | 3.42 |
| Parahippocampal gyrus, R[a] | 14 | -21 | -19 | 5 | 3.41 |
| Inferior frontal gyrus, triangular part, L | -38 | 22 | 5 | 6 | 3.40 |
| Middle cingulate & paracingulate gyri, L[a] | -6 | -16 | 24 | 5 | 3.39 |
| Lobule X of vermis[a] | 2 | -37 | -40 | 7 | 3.39 |
| Superior frontal gyrus, dorsolateral, R[a] | 24 | 50 | -5 | 8 | 3.39 |
| Lobule VI of cerebellar hemisphere, R | 35 | -45 | -27 | 5 | 3.39 |
| Superior temporal gyrus, L | -58 | -3 | -6 | 5 | 3.36 |
| Superior temporal gyrus, L | -53 | -6 | -10 | 6 | 3.31 |
| Postcentral gyrus, L | -61 | -2 | 21 | 8 | 3.31 |
| Hippocampus, R[a] | 24 | -35 | 14 | 5 | 3.30 |
| Middle occipital gyrus, R | 42 | -77 | 13 | 12 | 3.29 |
| Fusiform gyrus, L | -27 | -64 | -10 | 7 | 3.21 |
| Middle frontal gyrus, R | 35 | 30 | 29 | 6 | 3.19 |
| Superior frontal gyrus, dorsolateral, R | 32 | 66 | 2 | 5 | 3.19 |
| Inferior occipital gyrus, R | 42 | -80 | -5 | 8 | 3.16 |
| Temporal pole: superior temporal gyrus, R | 64 | 2 | 0 | 5 | 3.13 |
| Thalamus, pulvinar medial, R[a] | 10 | -22 | 18 | 7 | 3.03 |
| **UP < ER** | | | | | |
| Superior frontal gyrus, medial, L | -2 | 54 | 6 | 15175 | 10.05 |

| | | | | | |
|---|---|---|---|---|---|
| Precuneus, L | 2 | -61 | 27 | 20719 | 9.85 |
| Anterior orbital gyrus, L | -32 | 37 | -14 | 4091 | 8.90 |
| IFG pars orbitalis, R | 32 | 32 | -13 | 541 | 8.59 |
| Middle occipital gyrus, L | -38 | -66 | 24 | 1803 | 7.68 |
| Crus I of cerebellar hemisphere, R | 32 | -77 | -35 | 1288 | 7.40 |
| Insula, R | 42 | -11 | -3 | 3051 | 7.26 |
| Crus II of cerebellar hemisphere, L | -42 | -69 | -38 | 502 | 5.75 |
| Angular gyrus, R | 43 | -56 | 21 | 621 | 5.73 |
| Lobule VI of cerebellar hemisphere, L | -10 | -64 | -16 | 532 | 5.73 |
| Lingual gyrus, R | 10 | -69 | -8 | 154 | 5.72 |
| Middle temporal gyrus, L | -59 | -2 | -24 | 546 | 5.67 |
| Insula, L[a] | -29 | -30 | 24 | 35 | 5.16 |
| Inferior temporal gyrus, L | -59 | -58 | -10 | 154 | 5.10 |
| Insula, R[a] | 30 | -32 | 26 | 65 | 4.97 |
| Caudate nucleus, L[a] | -16 | 16 | 26 | 60 | 4.91 |
| Insula, L[a] | -22 | 26 | 11 | 21 | 4.73 |
| Caudate nucleus, R | 21 | -22 | 21 | 9 | 4.71 |
| Lobule IX of cerebellar hemisphere, R | 6 | -50 | -45 | 110 | 4.67 |
| Caudate nucleus, L | -16 | 10 | 22 | 20 | 4.66 |
| Middle cingulate & paracingulate gyri, L[a] | -14 | -16 | 34 | 36 | 4.66 |
| Hippocampus, L | -29 | -34 | -6 | 13 | 4.63 |
| Middle frontal gyrus, R | 48 | 42 | 6 | 273 | 4.61 |
| Caudate nucleus, L[a] | -14 | -3 | 29 | 63 | 4.59 |
| Olfactory cortex, R[a] | 6 | 8 | -16 | 8 | 4.49 |
| Lobule IV, V of cerebellar hemisphere, L | -10 | -46 | -18 | 35 | 4.46 |
| Lingual gyrus, R | 21 | -77 | -2 | 6 | 4.44 |
| Caudate nucleus, R[a] | 24 | -2 | 27 | 107 | 4.40 |
| Thalamus, pulvinar medial, L[a] | -21 | -27 | 18 | 42 | 4.35 |
| Gyrus rectus, R | 13 | 45 | -16 | 8 | 4.34 |
| Inferior frontal gyrus, triangular part, L | -46 | 29 | 3 | 193 | 4.32 |
| Lobule VIII of cerebellar hemisphere, L | -26 | -48 | -48 | 122 | 4.25 |
| Lobule VIII of vermis | 2 | -70 | -34 | 46 | 4.21 |
| Parahippocampal gyrus, R | 21 | -13 | -30 | 19 | 4.17 |
| Postcentral gyrus, L[a] | -30 | -19 | 32 | 51 | 4.17 |
| Middle temporal gyrus, L[a] | -40 | -45 | -3 | 18 | 4.13 |
| Lobule IV, V of cerebellar hemisphere, R | 18 | -56 | -14 | 31 | 4.12 |
| Middle cingulate & paracingulate gyri, L[a] | -22 | -30 | 38 | 18 | 4.12 |
| Temporal pole: middle temporal gyrus, L[a] | -38 | 11 | -45 | 11 | 4.08 |
| Inferior temporal gyrus, R | 53 | -37 | -22 | 34 | 4.08 |
| Middle temporal gyrus, R | 61 | -14 | -19 | 94 | 4.08 |
| Lobule IV, V of cerebellar hemisphere, L | -22 | -27 | -32 | 6 | 4.05 |
| Superior temporal gyrus, L | -48 | -32 | 22 | 63 | 4.05 |
| Supplementary motor area, L | -8 | -21 | 59 | 13 | 4.05 |
| Lobule IX of cerebellar hemisphere, L[a] | -19 | -46 | -37 | 11 | 4.04 |
| Inferior temporal gyrus, L | -59 | -30 | -24 | 12 | 4.02 |
| Middle cingulate & paracingulate gyri, L | -14 | -45 | 37 | 7 | 4.01 |

| | | | | | |
|---|---|---|---|---|---|
| Middle cingulate & paracingulate gyri, R | 3 | 8 | 35 | 33 | 4.00 |
| Anterior cingulate cortex, supracallosal, R[a] | 14 | 19 | 21 | 9 | 3.98 |
| Lobule IX of cerebellar hemisphere, L | -10 | -61 | -42 | 11 | 3.98 |
| Middle temporal gyrus, L | -61 | -43 | -11 | 45 | 3.95 |
| Inferior temporal gyrus, L | -56 | -14 | -30 | 28 | 3.91 |
| Precuneus, L[a] | 0 | -80 | 53 | 25 | 3.89 |
| Angular gyrus, L[a] | -24 | -45 | 30 | 11 | 3.88 |
| Hippocampus, R | 21 | -10 | -14 | 8 | 3.87 |
| Middle cingulate & paracingulate gyri, R[a] | 18 | -35 | 27 | 11 | 3.86 |
| Middle frontal gyrus, R[a] | 32 | 37 | 2 | 13 | 3.83 |
| Crus II of cerebellar hemisphere, L | -14 | -90 | -30 | 71 | 3.83 |
| Middle temporal gyrus, L | -67 | -27 | -16 | 27 | 3.81 |
| Unknown[a] | -26 | -40 | 27 | 19 | 3.79 |
| Supramarginal gyrus, L | -58 | -29 | 21 | 36 | 3.77 |
| Gyrus rectus, L[a] | 2 | 11 | -27 | 40 | 3.76 |
| Lingual gyrus, L[a] | -29 | -67 | 0 | 7 | 3.75 |
| Inferior temporal gyrus, L | -37 | 8 | -37 | 9 | 3.74 |
| Postcentral gyrus, L | -37 | -18 | 43 | 11 | 3.74 |
| Hippocampus, R[a] | 35 | -24 | -6 | 41 | 3.73 |
| Middle cingulate & paracingulate gyri, L[a] | -14 | -6 | 45 | 7 | 3.72 |
| Caudate nucleus, R | 21 | 5 | 24 | 21 | 3.72 |
| Hippocampus, R | 22 | -30 | -11 | 5 | 3.71 |
| Fusiform gyrus, R | 32 | -5 | -40 | 7 | 3.70 |
| Postcentral gyrus, L[a] | -35 | -14 | 35 | 8 | 3.68 |
| Crus I of cerebellar hemisphere, R | 11 | -90 | -24 | 8 | 3.68 |
| Anterior cingulate cortex, supracallosal, L | -5 | 16 | 26 | 12 | 3.68 |
| Lingual gyrus, R[a] | 14 | -94 | -19 | 11 | 3.67 |
| Precuneus, R[a] | 27 | -50 | 29 | 29 | 3.66 |
| Precuneus, L[a] | -19 | -50 | 40 | 13 | 3.65 |
| Caudate nucleus, L[a] | -21 | -35 | 24 | 8 | 3.65 |
| Inferior temporal gyrus, L | -43 | 11 | -40 | 8 | 3.65 |
| Caudate nucleus, L[a] | -22 | -8 | 21 | 13 | 3.64 |
| Insula, R[a] | 35 | -24 | 6 | 5 | 3.62 |
| Inferior parietal gyrus, excluding supramarginal and angular gyri, L[a] | -26 | -32 | 32 | 13 | 3.61 |
| Lobule X of cerebellar hemisphere, L[a] | -13 | -26 | -48 | 6 | 3.59 |
| Thalamus, ventral lateral, L | -10 | -10 | 16 | 10 | 3.58 |
| Insula, L[a] | -26 | 19 | 16 | 27 | 3.56 |
| Lobule IV, V of cerebellar hemisphere, R[a] | 22 | -35 | -30 | 9 | 3.56 |
| Middle cingulate & paracingulate gyri, L[a] | -16 | 10 | 34 | 6 | 3.55 |
| Middle frontal gyrus, R[a] | 29 | 13 | 35 | 6 | 3.53 |
| Heschl's gyrus, R[a] | 32 | -42 | 18 | 15 | 3.53 |
| Hippocampus, L[a] | -40 | -19 | -14 | 7 | 3.52 |
| Precuneus, L[a] | -16 | -53 | 32 | 14 | 3.51 |
| Middle occipital gyrus, L[a] | -21 | -82 | 5 | 18 | 3.50 |

| | | | | | |
|---|---|---|---|---|---|
| Middle cingulate & paracingulate gyri, R | 5 | 16 | 30 | 10 | 3.50 |
| Inferior temporal gyrus, R | 56 | -6 | -32 | 17 | 3.49 |
| Lobule IX of cerebellar hemisphere, R | 18 | -35 | -48 | 6 | 3.49 |
| Temporal pole: superior temporal gyrus, L | -32 | 19 | -27 | 14 | 3.48 |
| Temporal pole: middle temporal gyrus, R | 48 | 14 | -38 | 9 | 3.48 |
| Lobule IV, V of cerebellar hemisphere, L | -14 | -48 | -22 | 5 | 3.47 |
| Postcentral gyrus, R[a] | 38 | -29 | 32 | 6 | 3.46 |
| Lingual gyrus, R | 21 | -99 | -8 | 31 | 3.46 |
| Middle temporal gyrus, L | -53 | -24 | -13 | 17 | 3.46 |
| Middle occipital gyrus, L | -10 | -102 | 3 | 13 | 3.44 |
| Gyrus rectus, R | 19 | 19 | -14 | 6 | 3.42 |
| Hippocampus, R[a] | 37 | -43 | 0 | 11 | 3.40 |
| Middle occipital gyrus, L | -24 | -93 | 0 | 30 | 3.40 |
| Lobule IX of cerebellar hemisphere, R | 19 | -45 | -53 | 20 | 3.39 |
| Temporal pole: middle temporal gyrus, R | 29 | 21 | -40 | 5 | 3.38 |
| Temporal pole: superior temporal gyrus, L | -35 | 10 | -30 | 7 | 3.37 |
| Caudate nucleus, L[a] | -3 | 10 | 18 | 6 | 3.37 |
| Middle cingulate & paracingulate gyri, R[a] | 26 | -16 | 38 | 5 | 3.36 |
| Supramarginal gyrus, L | -62 | -32 | 30 | 49 | 3.36 |
| Superior parietal gyrus, R | 26 | -75 | 48 | 10 | 3.35 |
| Lingual gyrus, L | -14 | -78 | -3 | 5 | 3.34 |
| Parahippocampal gyrus, L | -19 | -22 | -26 | 8 | 3.32 |
| Caudate nucleus, R | 21 | 22 | 14 | 5 | 3.32 |
| Middle temporal gyrus, R[a] | 37 | -64 | 5 | 6 | 3.32 |
| Inferior temporal gyrus, R | 58 | -46 | -18 | 5 | 3.31 |
| Lobule IV, V of cerebellar hemisphere, R | 13 | -45 | -19 | 6 | 3.31 |
| Lenticular nucleus, putamen, L | -27 | 8 | 10 | 6 | 3.30 |
| Lobule VI of cerebellar hemisphere, R[a] | 26 | -45 | -34 | 8 | 3.30 |
| Lobule VIII of cerebellar hemisphere, L | -43 | -48 | -53 | 10 | 3.30 |
| Middle occipital gyrus, L[a] | -30 | -74 | 8 | 15 | 3.29 |
| Inferior temporal gyrus, L | -59 | -51 | -22 | 13 | 3.29 |
| Inferior temporal gyrus, L | -53 | -6 | -43 | 5 | 3.29 |
| Calcarine fissure and surrounding cortex, L | -26 | -61 | 11 | 5 | 3.29 |
| Middle cingulate & paracingulate gyri, L[a] | -18 | -24 | 37 | 9 | 3.29 |
| Caudate nucleus, R | 19 | -16 | 22 | 6 | 3.29 |
| Calcarine fissure and surrounding cortex, L | 3 | -99 | 0 | 7 | 3.28 |
| Gyrus rectus, R[a] | 8 | 19 | -30 | 9 | 3.27 |
| Inferior temporal gyrus, R | 51 | -51 | -27 | 5 | 3.27 |
| Lobule IX of cerebellar hemisphere, L[a] | -16 | -35 | -53 | 8 | 3.25 |
| Postcentral gyrus, R | 67 | -14 | 24 | 7 | 3.24 |
| Caudate nucleus, R[a] | 26 | -13 | 19 | 7 | 3.24 |
| Middle cingulate & paracingulate gyri, L[a] | -14 | -24 | 34 | 5 | 3.24 |
| Temporal pole: middle temporal gyrus, L | -27 | 10 | -43 | 6 | 3.24 |
| Lobule VIII of cerebellar hemisphere, R | 22 | -58 | -42 | 7 | 3.23 |
| Superior occipital gyrus, L | -19 | -82 | 43 | 6 | 3.21 |
| Olfactory cortex, L[a] | -5 | 6 | -21 | 5 | 3.21 |

| | | | | | |
|---|---|---|---|---|---|
| Inferior temporal gyrus, L | -53 | -43 | -16 | 5 | 3.21 |
| Inferior temporal gyrus, R | 56 | -54 | -22 | 5 | 3.21 |
| Crus I of cerebellar hemisphere, R | 21 | -90 | -21 | 6 | 3.21 |
| Calcarine fissure and surrounding cortex, L | -3 | -99 | -2 | 16 | 3.20 |
| Hippocampus, R[a] | 29 | -29 | 3 | 6 | 3.18 |
| Fusiform gyrus, R[a] | 43 | -35 | -11 | 7 | 3.17 |
| Insula, R | 30 | -21 | 18 | 8 | 3.17 |
| Supplementary motor area, R | 8 | -14 | 64 | 9 | 3.16 |
| Middle cingulate & paracingulate gyri, R[a] | 5 | -24 | 22 | 7 | 3.15 |
| Lobule VIII of vermis | -2 | -72 | -42 | 11 | 3.15 |
| Crus I of cerebellar hemisphere, L | -56 | -48 | -43 | 5 | 3.15 |
| Middle frontal gyrus, L | -38 | 40 | 22 | 9 | 3.12 |
| Superior frontal gyrus, medial, L | -3 | 58 | 42 | 9 | 3.11 |
| Precentral gyrus, R[a] | 27 | -13 | 35 | 7 | 3.11 |
| Supramarginal gyrus, L[a] | -38 | -40 | 26 | 5 | 3.11 |
| Supplementary motor area, R | 13 | -11 | 61 | 7 | 3.09 |
| Crus I of cerebellar hemisphere, R | 53 | -50 | -30 | 5 | 3.09 |
| Angular gyrus, L[a] | -29 | -51 | 21 | 6 | 3.09 |
| Inferior temporal gyrus, R | 61 | -10 | -30 | 6 | 3.09 |
| Lingual gyrus, R | 21 | -46 | -10 | 7 | 3.08 |
| Anterior cingulate cortex, supracallosal, R | 2 | 21 | 27 | 7 | 3.08 |
| Middle frontal gyrus, R | 48 | 50 | 11 | 10 | 3.07 |
| Caudate nucleus, R[a] | 27 | 5 | 19 | 5 | 3.07 |
| Inferior temporal gyrus, R | 58 | -30 | -21 | 6 | 3.07 |
| Hippocampus, L[a] | -35 | -8 | -16 | 7 | 3.03 |
| Lobule VIII of cerebellar hemisphere, R[a] | 29 | -46 | -42 | 5 | 3.02 |
| Insula, R[a] | 32 | 8 | 18 | 6 | 3.01 |
| Lingual gyrus, L | -27 | -93 | -18 | 11 | 3.00 |
| Crus II of cerebellar hemisphere, R | 46 | -45 | -42 | 9 | 2.99 |
| Lingual gyrus, L | -8 | -80 | -5 | 9 | 2.98 |
| Gyrus rectus, L | -6 | 50 | -19 | 6 | 2.97 |
| Anterior cingulate cortex, supracallosal, R[a] | 18 | 30 | 19 | 8 | 2.95 |
| Inferior temporal gyrus, R | 54 | 3 | -37 | 5 | 2.91 |
| Crus I of cerebellar hemisphere, L | -26 | -86 | -30 | 5 | 2.89 |
| Inferior occipital gyrus, L | -29 | -83 | -8 | 5 | 2.85 |
| Precentral gyrus, R | 22 | -16 | 61 | 6 | 2.80 |

Anatomic regions are defined using the Automated Anatomical Labelling Atlas 3 (AAL3; local maxima labelling, 1mm voxel edge)[1]. Significant clusters are thresholded at $P_{FDR} < .05$, $K_E \geq 5$ voxels. *T* values represent peak activation within the cluster.

[a]Cluster peak coordinates fall outside the defined regions, hence, were labelled using the nearest available anatomic region using AAL3. Regions labelled as 'Unknown' if no applicable AAL3 label can be defined.

*MNI*, Montreal Neurological Institute; *UP*, unexpected punishment; *ER*, expected reward; *L*, left; *R*, right.

**Supplementary Table 2. Significant perceptual decision uncertainty task-induced activation.**

| | MNI coordinates | | | Statistics | |
|---|---|---|---|---|---|
| | x | y | z | $K_E$ | T value |
| **IncorrFB > CorrFB** | | | | | |
| Supplementary motor area, L | 2 | 22 | 53 | 7104 | 10.27 |
| Insula, L | -32 | 21 | -3 | 5571 | 9.97 |
| Insula, R | 35 | 19 | -2 | 8309 | 9.62 |
| Superior temporal gyrus, R | 46 | -29 | -2 | 4456 | 7.44 |
| Middle temporal gyrus, L | -51 | -51 | 10 | 2479 | 6.79 |
| Thalamus, pulvinar medial, R[a] | 2 | -27 | 2 | 1963 | 6.61 |
| Middle frontal gyrus, L | -34 | 54 | 18 | 520 | 6.52 |
| Lobule IX of vermis | 0 | -53 | -38 | 72 | 6.49 |
| Middle temporal gyrus, L | -50 | -30 | -3 | 181 | 6.44 |
| Superior temporal gyrus, L | -45 | -27 | 10 | 312 | 5.92 |
| Lobule VI of cerebellar hemisphere, L | -30 | -58 | -29 | 251 | 5.81 |
| Red nucleus, L[a] | -8 | -27 | -13 | 35 | 5.76 |
| Red nucleus, R[a] | 2 | -21 | -21 | 96 | 5.62 |
| Insula, R | 43 | -3 | -14 | 65 | 5.54 |
| Crus I of cerebellar hemisphere, L | -19 | -70 | -30 | 247 | 5.13 |
| Posterior cingulate gyrus, L | -2 | -34 | 29 | 21 | 4.85 |
| Middle cingulate & paracingulate gyri, R | 8 | -26 | 29 | 136 | 4.84 |
| Lobule IX of cerebellar hemisphere, R[a] | 11 | -43 | -32 | 7 | 4.56 |
| Temporal pole: superior temporal gyrus, L | -42 | 11 | -24 | 24 | 4.54 |
| Precuneus, R | 11 | -69 | 54 | 281 | 4.53 |
| Calcarine fissure and surrounding cortex, L | -3 | -70 | 11 | 67 | 4.53 |
| Lobule IV, V of vermis[a] | 2 | -48 | -24 | 19 | 4.45 |
| Precuneus, L | -5 | -53 | 46 | 16 | 4.43 |
| Thalamus, lateral geniculate, R | 26 | -22 | -8 | 21 | 4.31 |
| Temporal pole: superior temporal gyrus, R | 37 | 19 | -27 | 5 | 4.30 |
| Calcarine fissure and surrounding cortex, L | -14 | -62 | 6 | 34 | 4.27 |
| Superior frontal gyrus, dorsolateral, L | -14 | 8 | 67 | 44 | 4.26 |
| Middle frontal gyrus, L | -29 | 51 | -11 | 9 | 4.21 |
| Lobule IV, V of vermis | 5 | -59 | -6 | 24 | 4.20 |
| Red nucleus, Ra | 3 | -16 | -8 | 25 | 4.20 |
| Parahippocampal gyrus, L[a] | -6 | -26 | -21 | 10 | 4.20 |
| Inferior temporal gyrus, L | -43 | -54 | -11 | 25 | 4.13 |
| Lingual gyrus, R | 14 | -61 | 8 | 156 | 4.10 |
| Superior temporal gyrus, L | -54 | -3 | -6 | 6 | 4.00 |
| Insula, R[a] | 38 | -8 | -10 | 5 | 3.94 |
| Inferior temporal gyrus, R | 50 | 2 | -40 | 8 | 3.93 |
| Thalamus, lateral geniculate, L | 19 | -26 | -6 | 21 | 3.93 |
| Medial orbital gyrus, R[a] | 13 | 38 | -30 | 17 | 3.91 |
| Superior frontal gyrus, dorsolateral, R | 14 | 58 | 27 | 28 | 3.90 |
| Inferior frontal gyrus, opercular part, L | -61 | 10 | 21 | 17 | 3.89 |
| Middle temporal gyrus, R | 69 | -22 | -14 | 15 | 3.87 |
| Middle temporal gyrus, R | 56 | -26 | -13 | 24 | 3.86 |
| Precuneus, L | -10 | -74 | 62 | 8 | 3.86 |

| | | | | | |
|---|---|---|---|---|---|
| Middle temporal gyrus, L | -53 | 8 | -29 | 9 | 3.85 |
| Fusiform gyrus, R | 32 | -48 | -21 | 12 | 3.84 |
| Supramarginal gyrus, L | -56 | -29 | 22 | 23 | 3.83 |
| Substantia nigra, pars reticulata, L[a] | -5 | -8 | -8 | 9 | 3.82 |
| Thalamus, pulvinar medial, L[a] | -11 | -30 | 16 | 8 | 3.82 |
| IFG pars orbitalis, L[a] | -51 | 45 | -8 | 5 | 3.79 |
| Middle temporal gyrus, R | 54 | -53 | -3 | 14 | 3.75 |
| Lobule III of cerebellar hemisphere, R[a] | 11 | -30 | -26 | 16 | 3.75 |
| Lobule I, II of vermis[a] | 3 | -37 | -29 | 5 | 3.74 |
| Medial orbital gyrus, R | 16 | 58 | -16 | 14 | 3.73 |
| Parahippocampal gyrus, L | -24 | -2 | -37 | 8 | 3.70 |
| Temporal pole: superior temporal gyrus, L | -54 | 16 | -22 | 14 | 3.68 |
| Precentral gyrus, L | -54 | 2 | 18 | 17 | 3.67 |
| Anterior cingulate cortex, supracallosal, R | 14 | 34 | 8 | 15 | 3.65 |
| Superior temporal gyrus, L | -59 | -8 | 5 | 5 | 3.65 |
| Middle temporal gyrus, L | -50 | 2 | -19 | 13 | 3.60 |
| Lobule IV, V of vermis | 2 | -53 | -8 | 26 | 3.60 |
| Middle frontal gyrus, L[a] | -53 | 18 | 42 | 7 | 3.60 |
| Inferior parietal gyrus, excluding supramarginal and angular gyri, L[a] | -56 | -37 | 56 | 9 | 3.59 |
| Lobule III of vermis | 3 | -42 | -14 | 10 | 3.59 |
| Lingual gyrus, L | -14 | -48 | -6 | 11 | 3.59 |
| Calcarine fissure and surrounding cortex, R | 19 | -67 | 10 | 7 | 3.56 |
| Supramarginal gyrus, R | 58 | -34 | 40 | 12 | 3.52 |
| Lobule VIIB of cerebellar hemisphere, L | -38 | -56 | -46 | 25 | 3.51 |
| Precentral gyrus, R[a] | 37 | -2 | 40 | 7 | 3.50 |
| Fusiform gyrus, R | 37 | -42 | -22 | 13 | 3.49 |
| Middle temporal gyrus, L | -46 | -3 | -30 | 7 | 3.49 |
| Precuneus, R | 8 | -54 | 48 | 6 | 3.48 |
| Lobule VI of cerebellar hemisphere, R | 11 | -69 | -27 | 5 | 3.47 |
| Middle temporal gyrus, L | -67 | -38 | 0 | 14 | 3.43 |
| Precuneus, R | 3 | -53 | 46 | 10 | 3.43 |
| Gyrus rectus, R | 8 | 50 | -19 | 5 | 3.41 |
| Supramarginal gyrus, L[a] | -42 | -40 | 26 | 5 | 3.41 |
| Insula, L | -35 | 3 | -2 | 6 | 3.41 |
| Lobule IV, V of vermis | 2 | -53 | -14 | 10 | 3.40 |
| Superior temporal gyrus, R | 62 | -8 | -8 | 5 | 3.39 |
| Middle temporal gyrus, L | -48 | -19 | -8 | 6 | 3.39 |
| Temporal pole: middle temporal gyrus, R | 48 | 11 | -34 | 8 | 3.37 |
| Parahippocampal gyrus, La | -6 | -2 | -32 | 6 | 3.37 |
| Temporal pole: superior temporal gyrus, R | 51 | 22 | -19 | 8 | 3.36 |
| Posterior orbital gyrus, R | 30 | 26 | -26 | 6 | 3.35 |
| IFG pars orbitalis, R | 40 | 42 | -8 | 6 | 3.34 |
| Rolandic operculum, L | -37 | -32 | 19 | 5 | 3.33 |
| Lobule VI of cerebellar hemisphere, R | 24 | -72 | -27 | 9 | 3.32 |
| Precuneus, L | -8 | -64 | 40 | 7 | 3.32 |
| Superior temporal gyrus, L | -58 | -27 | 3 | 5 | 3.31 |
| Lobule VIII of cerebellar hemisphere, R | 26 | -66 | -51 | 11 | 3.30 |
| Crus II of cerebellar hemisphere, L | -2 | -82 | -40 | 7 | 3.30 |

| | | | | | |
|---|---|---|---|---|---|
| Superior frontal gyrus, dorsolateral, L | -30 | 53 | -5 | 9 | 3.29 |
| Insula, R | 38 | 5 | -2 | 5 | 3.21 |
| Precuneus, R | 13 | -51 | 40 | 7 | 3.16 |
| Lobule VI of cerebellar hemisphere, L | -5 | -67 | -22 | 6 | 3.15 |
| Lobule IX of cerebellar hemisphere, L[a] | -11 | -48 | -32 | 5 | 3.14 |
| Inferior temporal gyrus, L | -46 | 6 | -40 | 7 | 3.11 |
| Inferior frontal gyrus, triangular part, R | 51 | 37 | -2 | 8 | 3.07 |
| Inferior parietal gyrus, excluding supramarginal and angular gyri, L | -54 | -56 | 38 | 9 | 2.99 |
| Inferior parietal gyrus, excluding supramarginal and angular gyri, R | 58 | -34 | 54 | 6 | 2.90 |
| **IncorrFB < CorrFB** | | | | | |
| Superior frontal gyrus, medial orbital, L | 0 | 50 | -5 | 19831 | 8.60 |
| Middle occipital gyrus, R | 29 | -78 | 16 | 9129 | 6.92 |
| Caudate nucleus, L[a] | -19 | 11 | 22 | 733 | 6.38 |
| Lenticular nucleus, putamen, L | -22 | 5 | -6 | 760 | 6.10 |
| Middle frontal gyrus, L | -32 | 30 | 48 | 1634 | 5.95 |
| Superior frontal gyrus, dorsolateral, R | 21 | 35 | 40 | 564 | 5.57 |
| Calcarine fissure and surrounding cortex, R[a] | 34 | -46 | 8 | 246 | 5.53 |
| Hippocampus, R | 35 | -19 | -19 | 309 | 5.28 |
| Inferior temporal gyrus, L | -56 | -26 | -19 | 16 | 5.28 |
| Hippocampus, L | -24 | -13 | -16 | 280 | 5.28 |
| Middle cingulate & paracingulate gyri, R | 11 | -2 | 40 | 74 | 5.13 |
| Parahippocampal gyrus, L | -27 | -38 | -8 | 197 | 5.10 |
| Hippocampus, R[a] | 30 | -30 | 6 | 106 | 5.04 |
| Inferior temporal gyrus, L | -58 | -58 | -11 | 189 | 5.03 |
| Lobule VIII of cerebellar hemisphere, L | -24 | -51 | -48 | 43 | 5.00 |
| Hippocampus, L | -24 | -37 | 0 | 57 | 4.98 |
| Lenticular nucleus, putamen, L[a] | -24 | -10 | 18 | 41 | 4.92 |
| Fusiform gyrus, R[a] | 34 | -56 | -2 | 54 | 4.89 |
| Precentral gyrus, L[a] | -22 | -24 | 54 | 52 | 4.89 |
| Lobule VIII of cerebellar hemisphere, R | 29 | -40 | -50 | 37 | 4.88 |
| Lingual gyrus, R | 19 | -58 | -13 | 53 | 4.70 |
| Inferior frontal gyrus, opercular part, R[a] | 35 | -2 | 26 | 13 | 4.70 |
| Supplementary motor area, L[a] | -11 | -18 | 56 | 20 | 4.66 |
| Lobule IX of cerebellar hemisphere, L | -13 | -48 | -56 | 100 | 4.65 |
| Cuneus, L | 2 | -77 | 27 | 59 | 4.58 |
| Thalamus, pulvinar medial, L[a] | -21 | -29 | 18 | 8 | 4.57 |
| Superior occipital gyrus, R | 24 | -64 | 42 | 54 | 4.54 |
| Fusiform gyrus, L | -21 | -43 | -19 | 50 | 4.51 |
| Lobule VIII of cerebellar hemisphere, L | -26 | -38 | -48 | 106 | 4.49 |
| Lobule IV, V of cerebellar hemisphere, R | 24 | -34 | -22 | 26 | 4.48 |
| Lobule IX of cerebellar hemisphere, R | 13 | -43 | -48 | 16 | 4.46 |
| Lobule IX of cerebellar hemisphere, L | -16 | -46 | -50 | 11 | 4.42 |
| Inferior temporal gyrus, L[a] | -61 | -30 | -29 | 11 | 4.41 |
| Crus I of cerebellar hemisphere, R | 42 | -70 | -32 | 56 | 4.38 |

| | | | | | |
|---|---|---|---|---|---|
| Crus I of cerebellar hemisphere, L | -48 | -50 | -30 | 12 | 4.35 |
| Middle temporal gyrus, L[a] | -34 | -48 | 10 | 14 | 4.35 |
| Fusiform gyrus, L | -26 | -54 | -14 | 19 | 4.25 |
| Anterior cingulate cortex, pregenual, R[a] | 19 | 43 | 10 | 9 | 4.22 |
| Superior frontal gyrus, dorsolateral, L[a] | -18 | 43 | 10 | 31 | 4.19 |
| Crus II of cerebellar hemisphere, R | 16 | -85 | -35 | 20 | 4.18 |
| Fusiform gyrus, L | -29 | -29 | -24 | 55 | 4.13 |
| Inferior temporal gyrus, L[a] | -38 | 5 | -46 | 5 | 4.12 |
| Superior frontal gyrus, dorsolateral, L | -24 | 2 | 67 | 17 | 4.12 |
| Rolandic operculum, L[a] | -40 | -19 | 26 | 24 | 4.11 |
| Precentral gyrus, R[a] | 27 | -10 | 48 | 25 | 4.11 |
| Angular gyrus, R | 45 | -67 | 32 | 112 | 4.10 |
| Superior frontal gyrus, dorsolateral, R | 24 | 32 | 29 | 10 | 4.09 |
| Caudate nucleus, L[a] | -8 | 6 | 22 | 40 | 4.07 |
| Hippocampus, R | 19 | -37 | 6 | 19 | 4.05 |
| Calcarine fissure and surrounding cortex, L[a] | -19 | -62 | 16 | 14 | 4.03 |
| Fusiform gyrus, L | -24 | -75 | -8 | 33 | 4.03 |
| Anterior cingulate cortex, supracallosal, L[a] | -18 | 29 | 18 | 17 | 4.01 |
| Precentral gyrus, R | 51 | 0 | 27 | 37 | 4.01 |
| IFG pars orbitalis, R | 35 | 34 | -11 | 33 | 4.00 |
| Superior temporal gyrus, R | 53 | -29 | 13 | 21 | 4.00 |
| Superior frontal gyrus, dorsolateral, R | 24 | 6 | 61 | 5 | 4.00 |
| Superior frontal gyrus, medial orbital, L[a] | -16 | 40 | -8 | 9 | 3.99 |
| Inferior temporal gyrus, L | -48 | -40 | -13 | 16 | 3.98 |
| Precentral gyrus, R[a] | 29 | -8 | 34 | 24 | 3.97 |
| Superior frontal gyrus, dorsolateral, L[a] | -18 | -10 | 51 | 13 | 3.96 |
| Rolandic operculum, L | -42 | -11 | 18 | 27 | 3.96 |
| Crus II of cerebellar hemisphere, L | -37 | -66 | -38 | 8 | 3.96 |
| Crus I of cerebellar hemisphere, R | 40 | -66 | -38 | 51 | 3.94 |
| Hippocampus, R[a] | 43 | -13 | -22 | 42 | 3.94 |
| Middle cingulate & paracingulate gyri, L | -6 | -3 | 45 | 10 | 3.94 |
| Crus I of cerebellar hemisphere, L | -40 | -80 | -22 | 37 | 3.93 |
| Middle cingulate & paracingulate gyri, L[a] | -13 | -19 | 48 | 13 | 3.93 |
| Supplementary motor area, L | -6 | -18 | 53 | 14 | 3.92 |
| Rolandic operculum, R | 62 | 8 | 10 | 12 | 3.90 |
| Inferior occipital gyrus, R | 35 | -78 | -11 | 11 | 3.87 |
| Hippocampus, L | -29 | -8 | -22 | 16 | 3.87 |
| Lobule VIII of cerebellar hemisphere, L | -29 | -51 | -54 | 7 | 3.86 |
| Parahippocampal gyrus, L[a] | -14 | -8 | -35 | 17 | 3.86 |
| Rolandic operculum, L | -43 | -6 | 11 | 8 | 3.85 |
| Insula, L[a] | -32 | -27 | 27 | 6 | 3.84 |
| Lobule VI of cerebellar hemisphere, L | -42 | -38 | -30 | 10 | 3.84 |
| Lenticular nucleus, putamen, L[a] | -22 | 13 | 11 | 7 | 3.83 |
| Insula, R | 40 | -8 | 22 | 7 | 3.83 |
| Insula, R | 38 | 3 | 6 | 17 | 3.82 |
| Lingual gyrus, L[a] | -29 | -59 | -3 | 9 | 3.82 |

| | | | | | |
|---|---|---|---|---|---|
| Middle temporal gyrus, L[a] | -43 | -21 | -14 | 11 | 3.82 |
| Middle frontal gyrus, L[a] | -27 | 29 | 26 | 5 | 3.81 |
| Unknown[a] | -3 | -19 | -45 | 14 | 3.81 |
| Fusiform gyrus, R[a] | 24 | -10 | -45 | 5 | 3.80 |
| Postcentral gyrus, L | -64 | -6 | 30 | 21 | 3.79 |
| Superior frontal gyrus, dorsolateral, R[a] | 21 | 32 | 24 | 6 | 3.78 |
| Crus I of cerebellar hemisphere, L | -40 | -70 | -32 | 8 | 3.77 |
| Inferior temporal gyrus, R | 50 | -74 | -11 | 8 | 3.76 |
| Middle cingulate & paracingulate gyri, L | -13 | -43 | 50 | 5 | 3.75 |
| Inferior temporal gyrus, L | -53 | -54 | -27 | 10 | 3.75 |
| Lingual gyrus, R | 26 | -64 | 0 | 6 | 3.74 |
| Lobule VI of cerebellar hemisphere, L | -11 | -61 | -14 | 26 | 3.73 |
| Middle cingulate & paracingulate gyri, R[a] | 18 | -6 | 42 | 16 | 3.73 |
| Inferior temporal gyrus, R | 43 | -6 | -37 | 10 | 3.72 |
| Inferior temporal gyrus, L | -50 | -48 | -22 | 5 | 3.71 |
| Lobule IX of cerebellar hemisphere, L[a] | -11 | -35 | -51 | 29 | 3.71 |
| Lenticular nucleus, putamen, L[a] | -19 | 22 | -8 | 8 | 3.71 |
| Calcarine fissure and surrounding cortex, L | -3 | -98 | 2 | 42 | 3.70 |
| Posterior orbital gyrus, L | -22 | 27 | -19 | 6 | 3.70 |
| Inferior temporal gyrus, L[a] | -43 | -40 | -10 | 21 | 3.70 |
| Fusiform gyrus, L | -29 | -48 | -14 | 7 | 3.69 |
| Parahippocampal gyrus, L | -29 | -27 | -18 | 15 | 3.69 |
| Lobule VI of cerebellar hemisphere, L | -22 | -67 | -16 | 7 | 3.68 |
| Postcentral gyrus, L[a] | -24 | -29 | 50 | 10 | 3.68 |
| Lobule VIII of cerebellar hemisphere, L | -26 | -62 | -40 | 9 | 3.67 |
| Superior frontal gyrus, dorsolateral, R | 18 | 35 | 30 | 8 | 3.66 |
| Paracentral lobule, L[a] | -14 | -27 | 58 | 6 | 3.66 |
| Hippocampus, L | -11 | -40 | 8 | 7 | 3.64 |
| Crus I of cerebellar hemisphere, L | -56 | -54 | -40 | 5 | 3.62 |
| Posterior orbital gyrus, R | 27 | 26 | -13 | 13 | 3.59 |
| Parahippocampal gyrus, R[a] | 19 | -18 | -32 | 11 | 3.59 |
| Crus I of cerebellar hemisphere, R | 13 | -88 | -21 | 5 | 3.59 |
| Lobule VIII of cerebellar hemisphere, R | 21 | -45 | -54 | 10 | 3.58 |
| Inferior temporal gyrus, R | 58 | -16 | -32 | 6 | 3.58 |
| Middle cingulate & paracingulate gyri, R[a] | 16 | 8 | 37 | 8 | 3.57 |
| Lobule IV, V of cerebellar hemisphere, L | -14 | -56 | -19 | 9 | 3.57 |
| Amygdala, R | 30 | 0 | -14 | 11 | 3.56 |
| Insula, R | 40 | -16 | 8 | 8 | 3.54 |
| Crus I of cerebellar hemisphere, R | 30 | -82 | -35 | 5 | 3.52 |
| Crus I of cerebellar hemisphere, R | 19 | -86 | -27 | 6 | 3.51 |
| Fusiform gyrus, R | 38 | -77 | -18 | 14 | 3.51 |
| Middle frontal gyrus, L | -38 | 35 | 27 | 27 | 3.50 |
| Caudate nucleus, L | -10 | 14 | 0 | 6 | 3.49 |
| Thalamus, pulvinar medial, R[a] | 5 | -26 | 16 | 7 | 3.48 |
| Middle occipital gyrus, L | -24 | -69 | 38 | 8 | 3.48 |
| Postcentral gyrus, R | 66 | 0 | 16 | 15 | 3.47 |

| | | | | | |
|---|---|---|---|---|---|
| Anterior cingulate cortex, supracallosal, R[a] | 6 | 30 | 8 | 5 | 3.47 |
| Crus I of cerebellar hemisphere, R | 32 | -74 | -35 | 11 | 3.44 |
| Rolandic operculum, R | 42 | -3 | 11 | 5 | 3.42 |
| Lobule IV, V of cerebellar hemisphere, L | -14 | -48 | -16 | 6 | 3.41 |
| Anterior cingulate cortex, subgenual, R[a] | 10 | 29 | -3 | 8 | 3.41 |
| Parahippocampal gyrus, R | 29 | -27 | -21 | 7 | 3.41 |
| Middle cingulate & paracingulate gyri, L | -13 | -38 | 45 | 5 | 3.40 |
| Lobule VI of cerebellar hemisphere, R | 34 | -72 | -22 | 6 | 3.38 |
| Precuneus, L | -14 | -58 | 16 | 6 | 3.36 |
| Crus I of cerebellar hemisphere, R | 45 | -46 | -32 | 11 | 3.36 |
| Middle cingulate & paracingulate gyri, L | -2 | -27 | 46 | 7 | 3.33 |
| Caudate nucleus, L[a] | -18 | -24 | 30 | 5 | 3.32 |
| Rolandic operculum, L[a] | -29 | -34 | 19 | 5 | 3.31 |
| Crus I of cerebellar hemisphere, R | 42 | -43 | -34 | 6 | 3.30 |
| Superior frontal gyrus, dorsolateral, L | -21 | 58 | 14 | 11 | 3.29 |
| Crus I of cerebellar hemisphere, R | 53 | -61 | -35 | 5 | 3.29 |
| Superior frontal gyrus, dorsolateral, L | -21 | 5 | 56 | 11 | 3.29 |
| Rolandic operculum, L[a] | -32 | -43 | 19 | 5 | 3.28 |
| Lenticular nucleus, putamen, L | -27 | -6 | -8 | 14 | 3.28 |
| Inferior temporal gyrus, L | -56 | -24 | -26 | 8 | 3.27 |
| Fusiform gyrus, R | 24 | -40 | -13 | 7 | 3.27 |
| Hippocampus, R[a] | 38 | -6 | -24 | 8 | 3.26 |
| Middle frontal gyrus, L | -40 | 37 | 32 | 5 | 3.26 |
| Cuneus, L[a] | -22 | -59 | 22 | 6 | 3.24 |
| Middle frontal gyrus, R | 29 | 38 | 27 | 8 | 3.24 |
| Inferior frontal gyrus, triangular part, L[a] | -51 | 42 | 13 | 6 | 3.24 |
| Unknown[a] | 6 | -24 | -45 | 14 | 3.22 |
| Fusiform gyrus, R[a] | 21 | 3 | -43 | 7 | 3.22 |
| Angular gyrus, L | -50 | -62 | 40 | 6 | 3.18 |
| Middle cingulate & paracingulate gyri, L | -3 | -6 | 37 | 6 | 3.18 |
| Lingual gyrus, L | -11 | -58 | -8 | 6 | 3.16 |
| Superior parietal gyrus, L | -16 | -56 | 45 | 6 | 3.16 |
| Fusiform gyrus, L[a] | -30 | -67 | -3 | 9 | 3.15 |
| Precuneus, R[a] | 29 | -58 | 19 | 5 | 3.14 |
| Lingual gyrus, R | 10 | -75 | -2 | 5 | 3.13 |
| Insula, L | -34 | -14 | 3 | 10 | 3.12 |
| Middle occipital gyrus, L | -30 | -82 | 29 | 6 | 3.11 |
| Inferior occipital gyrus, R | 38 | -74 | -13 | 7 | 3.11 |
| Precentral gyrus, L | -51 | -3 | 26 | 6 | 3.11 |
| Crus I of cerebellar hemisphere, L | -38 | -75 | -29 | 5 | 3.09 |
| Lobule IV, V of cerebellar hemisphere, L | -8 | -50 | -6 | 6 | 3.08 |
| Precuneus, R[a] | 19 | -46 | 19 | 5 | 3.07 |
| Inferior occipital gyrus, R | 38 | -78 | -5 | 7 | 3.05 |
| Cuneus, R | 18 | -98 | 8 | 5 | 3.01 |
| Middle frontal gyrus, L[a] | -22 | 13 | 34 | 7 | 3.00 |
| Calcarine fissure and surrounding cortex, R | 8 | -93 | 0 | 5 | 3.00 |

| | | | | | |
|---|---|---|---|---|---|
| Inferior temporal gyrus, R | 54 | -46 | -27 | 5 | 3.00 |
| Inferior parietal gyrus, excluding supramarginal and angular gyri, L | -30 | -77 | 45 | 5 | 2.94 |
| Middle frontal gyrus, L | -24 | 8 | 48 | 6 | 2.94 |

Anatomic regions are defined using the Automated Anatomical Labelling Atlas 3 (AAL3; local maxima labelling, 1mm voxel edge)[1]. Significant clusters are thresholded at $P_{FDR} < .05$, $K_E \geq 5$ voxels. *T* values represent peak activation within the cluster.

[a]Cluster peak coordinates fall outside the defined regions, hence, were labelled using the nearest available anatomic region using AAL3. Regions labelled as 'Unknown' if no applicable AAL3 label can be defined.

*MNI*, Montreal Neurological Institute; *IncorrFB*, incorrect feedback; *CorrFB*, correct feedback; *L*, left; *R*, right.

**Supplementary Table 3. Bayesian model-averaged DCM parameters for endogenous and modulatory connections.**

| | Reversal learning task | | | Perceptual decision uncertainty task | | |
|---|---|---|---|---|---|---|
| **Connection** | **Ep** | **Cp** | **Pp** | **Ep** | **Cp** | **Pp** |
| **Endogenous connections[a] (A-matrix)** | | | | | | |
| Hb → Hb | 0.09 | 0.0044 | .79 | -0.31 | 0.0035 | 1.00* |
| Hb → VTA | -0.14 | 0.0017 | 1.00* | 0.04 | 0.0026 | .51 |
| Hb → DRN | -0.57 | 0.0019 | 1.00* | 0.00 | 0.0000 | .00 |
| VTA → VTA | 0.00 | 0.0000 | .00 | -0.19 | 0.0043 | .98* |
| VTA → Hb | 0.64 | 0.0023 | 1.00* | 0.00 | 0.0000 | .00 |
| VTA → dACC | 0.11 | 0.0023 | .95 | 1.01 | 0.0033 | 1.00* |
| DRN → DRN | -0.30 | 0.0026 | 1.00* | -0.22 | 0.0040 | .99* |
| DRN → Hb | -0.14 | 0.0020 | .99* | -0.10 | 0.0046 | .82 |
| DRN → mPFC | 0.00 | 0.0000 | .00 | -0.93 | 0.0035 | 1.00* |
| dACC → dACC | 0.07 | 0.0042 | .69 | 0.00 | 0.0000 | .00 |
| dACC → Hb | -0.15 | 0.0026 | .99* | 0.00 | 0.0000 | .00 |
| dACC → VTA | -0.25 | 0.0020 | 1.00* | -0.24 | 0.0024 | 1.00* |
| mPFC → mPFC | -0.30 | 0.0023 | 1.00* | 0.05 | 0.0044 | .58 |
| mPFC → Hb | 0.00 | 0.0000 | .00 | 0.00 | 0.0000 | .00 |
| mPFC → DRN | -0.10 | 0.0015 | .95* | 0.28 | 0.0023 | 1.00* |
| **Modulatory connections[b] (B-matrix)** | | | | | | |
| **Negative feedback[c] (UP/IncorrFB)** | | | | | | |
| Hb → VTA | -3.65 | 0.1302 | 1.00* | -1.24 | 0.0398 | 1.00* |
| Hb → DRN | -1.61 | 0.1064 | 1.00* | -0.86 | 0.0322 | 1.00* |
| VTA → dACC | 3.26 | 0.0621 | 1.00* | -1.38 | 0.0504 | 1.00* |
| DRN → mPFC | 0.68 | 0.1080 | .90 | 1.49 | 0.0524 | 1.00* |
| dACC → Hb | 0.30 | 0.1170 | .56 | 0.71 | 0.0268 | 1.00* |
| **Positive feedback[d] (ER/CorrFB)** | | | | | | |
| Hb → VTA | 3.93 | 0.0763 | 1.00* | 2.70 | 0.0420 | 1.00* |
| Hb → DRN | 2.72 | 0.0531 | 1.00* | 1.16 | 0.0315 | 1.00* |
| VTA → dACC | 1.73 | 0.0565 | 1.00* | -0.44 | 0.0819 | .82 |
| DRN → mPFC | -9.33 | 0.1109 | 1.00* | 0.00 | 0.0000 | .00 |
| mPFC → Hb | 0.00 | 0.0000 | .00 | 0.00 | 0.0000 | .00 |

[a]Endogenous parameters reflect the average effective coupling between regions across experimental conditions (context-independent).

[b]Modulatory parameters reflect the changes in effective coupling between regions induced by feedback (content-dependent).

[c]Negative feedback is defined as unexpected punishment for the reversal learning task and incorrect response for the perceptual decision uncertainty task.

[d]Positive feedback refers to expected reward for the reversal learning task and correct response for the perceptual decision uncertainty task.

*Posterior probability (*Pp*) exceeding .95 provides sufficient evidence for a non-zero group effect [2].

*Ep*, posterior expectation; *Cp*, posterior covariance; *Pp*, posterior probability; *Hb,* Habenula; *VTA*, ventral tegmental area; *DRN*, dorsal raphe nucleus; *dACC*, dorsal anterior cingulate cortex; *mPFC*, medial prefrontal cortex; *UP*, unexpected punishment; *IncorrFB*, incorrect feedback; *ER*, expected reward; *CorrFB*, correct feedback.

**Supplementary Table 4. Demographic information and additional measures for the final samples.**

| | Reversal task (n = 47) | | Perceptual uncertainty task (n = 43) | |
|---|---|---|---|---|
| | **Male** | **Female** | **Male** | **Female** |
| **Sex** | 24 | 23 | 23 | 20 |

| | Reversal task (n = 47) | | | | | | | Perceptual uncertainty task (n = 43) | | | | | | |
|---|---|---|---|---|---|---|---|---|---|---|---|---|---|---|
| | **African** | **Asian** | **Australian (non-ATSI)** | **European** | **Middle Eastern** | **North American** | **South American** | **African** | **Asian** | **Australian (non-ATSI)** | **European** | **Middle Eastern** | **North American** | **South American** |
| **Ethno-cultural groups** | 1 | 30 | 11 | 1 | 0 | 2 | 2 | 0 | 28 | 9 | 2 | 1 | 2 | 1 |

| | Reversal task (n = 47) | | | | Perceptual uncertainty task (n = 43) | | | |
|---|---|---|---|---|---|---|---|---|
| | **Mean** | **SD** | **Minimum** | **Maximum** | **Mean** | **SD** | **Minimum** | **Maximum** |
| **Age (years)** | 28.9 | 8.7 | 18 | 59 | 29.4 | 8.9 | 18 | 59 |
| **QIDS-C[a] Total** | 2.0 | 1.7 | 0 | 6 | 2.0 | 1.7 | 0 | 6 |
| **SHAPS[b]** | 34.6 | 4.8 | 42 | 24 | 34.6 | 4.4 | 42 | 24 |

[a]Quick inventory of depressive symptomatology – clinician-rated. A 16-item semi-structured interview assessing the severity of the nine diagnostic symptom criteria of major depressive disorder used in the diagnostic and statistical manual of mental disorders 5th edition over the past 7 days. For each item, the scale varies from 0 to 3 where higher scores are indicative of depressive symptomatology[3].
[b]Snaith-Hamilton pleasure scale. A self-report measure consisting of 14 items that assess the level of anhedonia experienced by participants based on hedonic tone over the past few days. For each item, the scale varies from Strongly Disagree = 0 to Strongly Agree = 3[4].
*ATSI*, Aboriginal and/or Torres Strait Islander; *SD*, standard deviation.

**Supplementary Figures**

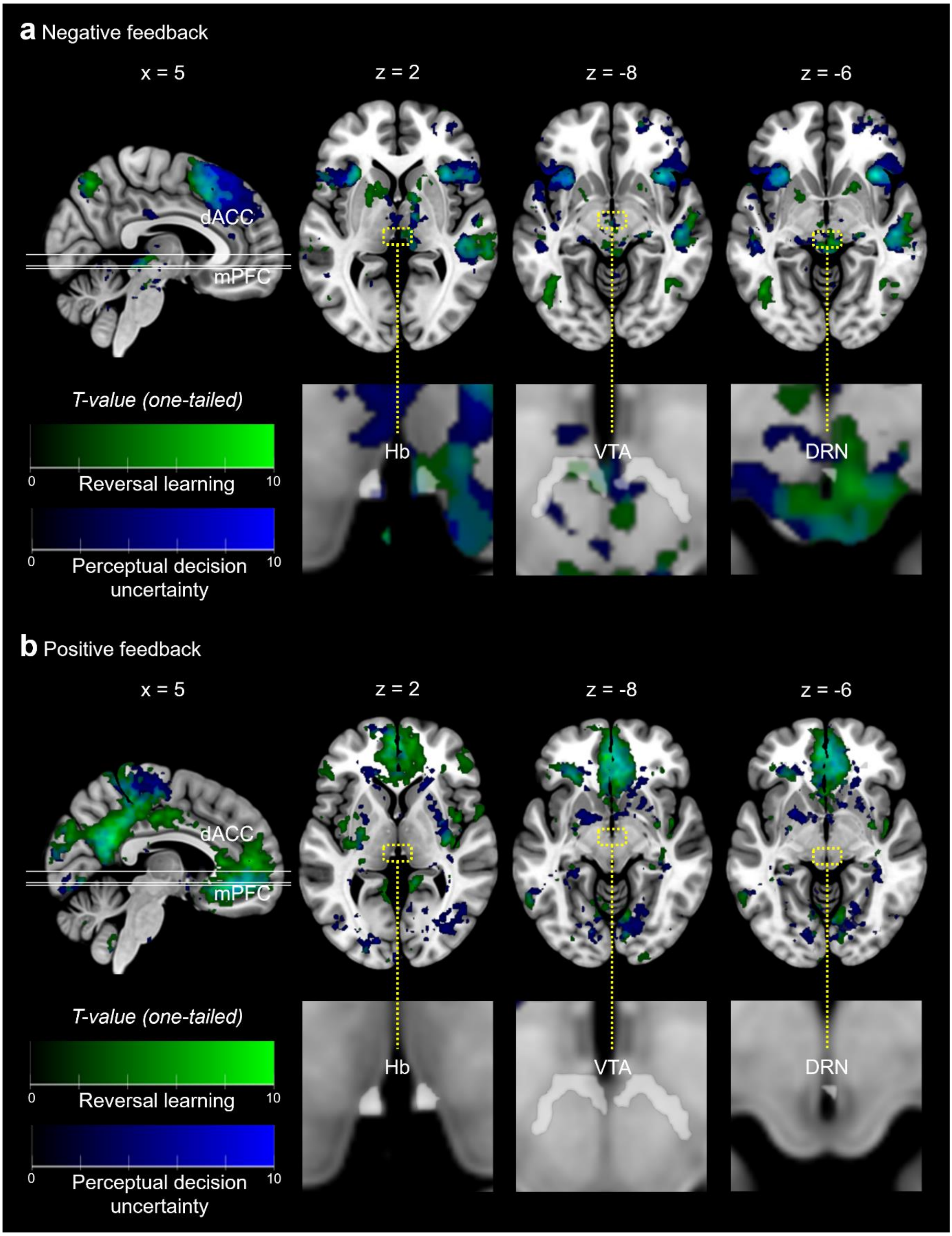

**Supplementary Figure 1. Overlapping activation for contrasts of interest across both the reversal learning and perceptual decision uncertainty tasks. a** Whole-brain GLM results assessing negative > positive feedback. **b** Whole-brain GLM results assessing negative < positive feedback. For both tasks, significant results ($P_{FDR} < .05$, $K_E \geq 5$) are presented on the MNI 152 template brain. The color maps represent corresponding *t*-statistic values for a one-tailed *t* test. Green colors correspond to regions of significant activation/deactivation during the reversal learning task (unexpected punishment; expected reward). Blue colors correspond to regions of significant activation/deactivation during the perceptual decision uncertainty task (incorrect response; correct response). Our targeted regions of interest are highlighted in their respective anatomical planes with translucent masks outlining the subcortical regions. *dACC*, dorsal anterior cingulate cortex; *MPFC*, medial prefrontal cortex; *Hb*, habenula; *VTA*, ventral tegmental area; *DRN*, dorsal raphe nucleus; *FDR*, false discovery rate; $K_E$, voxel cluster-extent threshold; *MNI,* Montreal Neurological Institute.

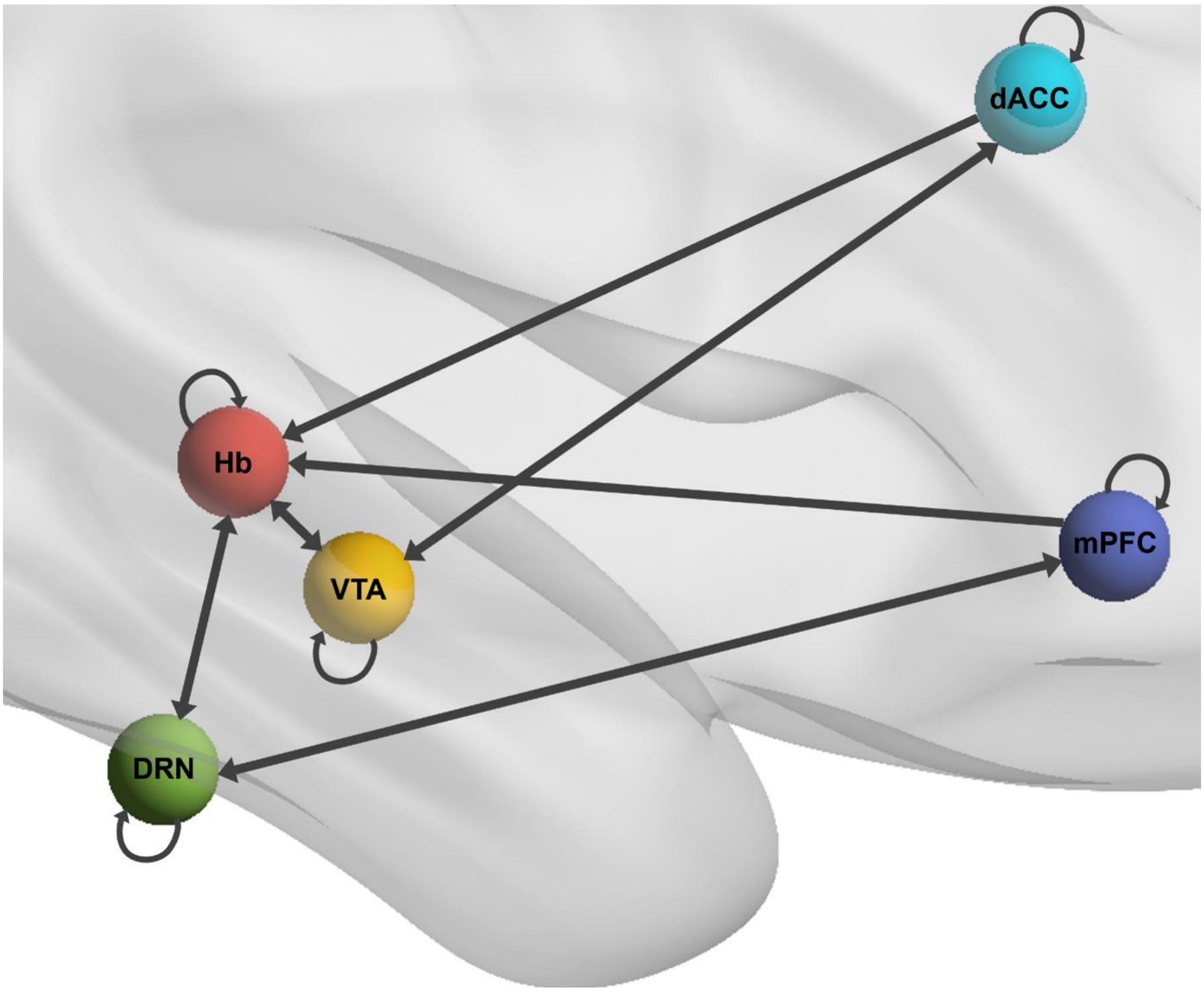

**Supplementary Figure 2. Endogenous DCM model space.** For both tasks, this DCM model was constructed and estimated for each subject. The model assumed bidirectional endogenous connections between most of the regions except for unidirectional endogenous connections from the cortical regions to the Hb. Endogenous self-inhibitory connections were also modelled. *Hb*, habenula; *VTA*, ventral tegmental area; *DRN*, dorsal raphe nucleus; *dACC*, dorsal anterior cingulate cortex; *mPFC*, medial prefrontal cortex.
